# Thermodynamic Profiling of Glucose Translocation in POPC embedded GLUT-4 protein models

**DOI:** 10.64898/2026.08.02.742329

**Authors:** S. Govinda, Suhotra Das, Allen Lobo, Chandrashekhar N. Rahul, Daniel Andrew Gideon

## Abstract

The facilitative transport of hexose through the GLUcose Transporter type 4 (GLUT4) is essential for cellular metabolism and is regulated by allosteric nucleotide interactions. In this study, we conducted a comparative biophysical analysis of glucose translocation using both the native Cryo-EM structure (7WSN) and the AI-predicted AlphaFold model using Steered Molecular Dynamics (SMD) at physiological (310.15 K) and attenuated (303.15 K) temperatures with two different allosteric modulators, ATP and ADP. While the AlphaFold model managed to capture baseline static topologies, dynamic profiling revealed that it intrinsically over-optimizes for static stability, immediately collapsing into a sterically occluded, hyper-packed artifact. Translocation through this constricted geometry forces severe steric solvent exclusion, abruptly stripping the substrate’s hydration shell and resulting in immediate and immense thermodynamic friction (>300 kJ/mol). Under ATP-bound physiological strain, this hyper-bonded structural clamp prevents functional relaxation, ultimately inducing a catastrophic kinematic failure. Furthermore, ensemble Dynamic Cross-Correlation Matrix (DCCM) analysis demonstrates that the AI-predicted model suffers from persistent rigid-body locking, forcing the transmembrane domain to behave as a kinetically trapped conformation. In contrast, the functionally hydrated Cryo-EM architecture actively maintains a dual-gating pore and successfully isolates the exact mechanical lever driving transport: a native Proline hinge (PRO379) that seamlessly routes allosteric signals from the nucleotide anchor to the pore gate. Within the AlphaFold model, this critical allosteric wiring is disrupted, forcing mechanical coupling through an unphysiological, hyper-correlated aromatic pathway. These findings yield novel atomistic insights into GLUT4 gating mechanics, while definitively establishing that rigid AI-generated architectures lack the essential internal free volume and conformational plasticity required to sustain accurate thermodynamic profiling of dynamic membrane transporters.

## 1. Introduction

D-Glucose, life’s primary metabolic currency, crosses the plasma membrane via the Solute Carrier 2 (SLC2) family of facilitative glucose transporters. Homeostatic regulation of blood glucose levels within the physiological range (70–140 mg/dL) depends on the GLUcose Transporter type 4 (GLUT4) protein, a multi-pass transmembrane protein highly expressed in insulin-responsive skeletal muscle and adipose tissues. Despite its primary sequence being determined in 1989, resolving its three-dimensional(3D) structure, which relies on nanoscale thermal fluctuations and conformational flexibility to achieve transport coordination, remained a bottleneck in structural biology. As a member of the Major Facilitator Superfamily (MFS), GLUT4 functions through an alternating-access mechanism requiring continuous transitions between inward-facing, outward-facing and occluded conformations. These transitions depend upon coordinated nanoscale thermal fluctuations, conformational flexibility and long-range allosteric communication throughout the transmembrane helices, making the protein inherently difficult to resolve experimentally. Consequently, for more than three decades, computational investigations relied predominantly on homology models derived from the structurally related GLUT1 transporter (Mohan S et al., 2009; Paul W. Hruz, 2001). Consequently, early *in silico* molecular dynamics simulations relied heavily on the GLUT 1 structural templates. A definitive breakthrough occurred in 2022 when the first high-resolution Cryo-EM structure of human GLUT4 captured in an inward-facing conformation, was discovered (Yuan et al., 2022), rapidly followed by computational structural predictions generated by DeepMind’s AlphaFold architecture (Jumper et al., 2021),with the predicted model made public in 2025. Both these models are shown in figure 1.

**Figure 1.**
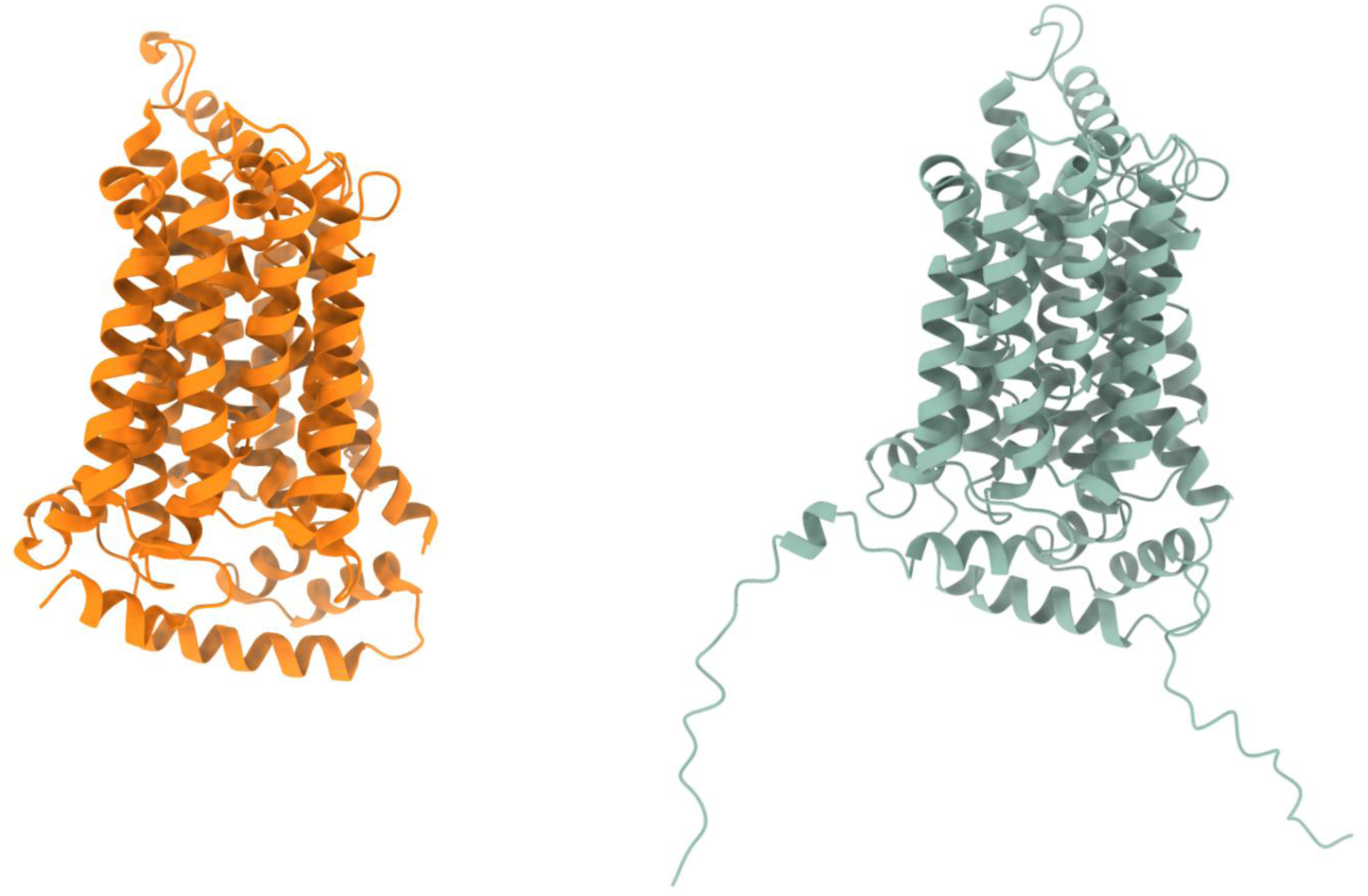
The structural models of GLUT4. Left Cryo-EM model. Right - AlphaFold predicted model.

Under basal conditions, GLUT4 resides within cytoplasmic vesicles (Drobiova et al., 2025) and translocates to the plasma membrane in response to extracellular insulin. Once embedded into the lipid bilayer, it facilitates downstream glucose diffusion through an alternating access mechanism (Figure 2). This macromolecular machinery, acts as a nanoscale valve, requiring the substrate (glucose) concentration gradient to navigate the channel’s electrostatic and Van der Waals potential energy barriers. The resulting accompanying conformational transitions dynamically alters the internal pore’s radius and electrostatic surface potential, efficiently driving substrate flux, all the while resisting backward Brownian motion. Elucidating these biophysical mechanisms remains critical for understanding the dysfunctions underlying various metabolic disorders globally.

**Figure 2.**
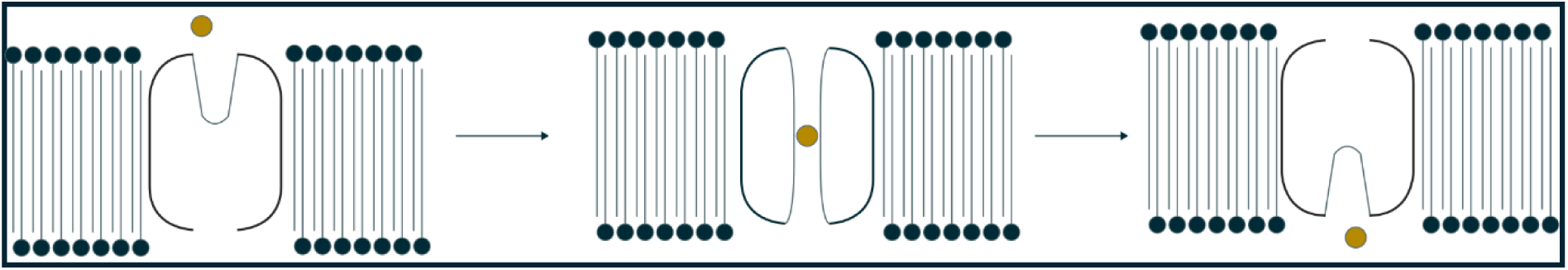
Representation of the change in channel conformation during transport.

Following transport into the cytoplasm, glucose undergoes rapid phosphorylation by hexokinase, as a part of the Glycolysis metabolic pathway (Lehninger et al., 2008), effectively preventing its diffusion back across the membrane, as phosphorylation imposes a steep energy barrier that ensures unidirectional transport while priming the molecule for cellular metabolism. ATP has been reported to interact with cytoplasmic regions of GLUT4 and homologous glucose transporters, altering local conformational geometry and modifying the electrostatic environment surrounding the transport pathway. Such interactions may increase the energetic cost of glucose translocation by modulating the free-energy landscape governing conformational transitions

This presence of ATP likely serves as an additional regulatory layer, occurring under two primary conditions:

- A diminished glucose gradient, where transport flux becomes slower than intracellular phosphorylation rate, leading to localized ATP crowding near the lower helices.
- Intracellular glucose approaches saturation within the cell, which increases the probability of stochastic ATP-protein encounters.

Accordingly, we hypothesize that nucleotide binding differentially remodels the structural, energetic and dynamic properties of GLUT4, thereby modulating the efficiency of glucose transport. Owing to differences in phosphate composition and charge distribution, ADP is anticipated to exert a substantially weaker allosteric influence than ATP.

The molecular basis of GLUT4 regulation has been investigated extensively over the past three decades. Historically, the molecular mechanisms of insulin-stimulated glucose uptake was first presented by the 1989 cDNA cloning of GLUT4 from skeletal muscle (Birnbaum, 1989). This seminal study revealed tissue-specific expression across adipose, skeletal, and cardiac tissues and established that insulin redistributes existing transporters rather than triggering *de novo* protein synthesis. Due to early experimental challenges with GLUT4, researchers often utilized the homologous GLUT1 isoform to map facilitative diffusion (Paul W. Hruz, 2001), successfully identifying the conserved <u>QQLS</u> glucose-binding motif (often extended to <u>LSQQLS</u> in computer simulations to ensure high-fidelity docking). The critical role of GLUT4 was subsequently validated by *in vivo* knockout mice models, which presented severely impaired insulin signalling and profound diabetic phenotypes (Huang & Czech, 2007; Kim et al., 2001). Furthermore, experimental investigations into nucleotide mediated metabolic cross-talk revealed several high-affinity ATP, ADP, and AMP binding sites, notably the conserved sequence <u>GRRTLHL</u> (Carruthers & Helgerson, 1989).

Aberrant regulation of GLUT4 surface expression and gating is directly implicated in severe metabolic pathologies, including Type 2 diabetes mellitus, neurodegenerative disorders such as Alzheimer’s disease, and malignant cell proliferation in highly glycolytic, proliferative cancers. Post-translational mechanistic studies indicate that strict de-phosphorylation networks govern vesicular translocation steps (Sadler et al., 2013). ATP interactions in homologous proteins exhibit concentration-dependent transport modulation (Cloherty et al., 2002), identifying allosteric binding domains sensitive to non-competitive small-molecule inhibitors like genistein (Bazuine et al., 2005). To probe these subtle molecular events, computational biophysics and Molecular Dynamics (MD) simulations (McCammon et al., 1977) have proven indispensable. Recent homology modelling (Mohan S et al., 2009) and corresponding docking studies indicate that ATP binding induces a highly compact, low-entropy structural state that effectively prevents functional channel opening and suppresses hexose transport (Bazuine et al., 2005; Mohan et al., 2010).

Despite recent milestones in capturing static structural views via Cryo-EM and deep-learning pipelines, a major knowledge gap persists regarding the non-equilibrium molecular mechanics of GLUT4 under active, non-idealized conditions. Current literature remains biased toward static interpretations of transporter topology, leaving unresolved how ambient thermal fluctuations, transmembrane vector stress, and allosteric nucleotide docking interact dynamically to alter local free energy barriers, water-wire networks, and transport equilibrium.

Concurrently, the rapid proliferation of AI-driven structural biology has catalysed a widespread reliance on AI-predicted models, notably the AlphaFold archive, for high-throughput computational drug screening and dynamic simulations. While these machine-learning architectures excel at predicting the idealized, global conformational minima of static, unliganded topologies, we hypothesize that they intrinsically over-optimize intramolecular contact networks to maximize static stability. This optimization occurs at the severe expense of the configurational entropy, internal free volume, and low-energy alternate states required to support functional allosteric gating. In highly dynamic membrane transport proteins such as GLUT4, this artifactually steric over-packing is hypothesized to collapse the water-filled pore lumen and suppress the independent helical flexibility required for the alternating access mechanism, effectively trapping the transporter in a hyper-bonded, dynamically arrested, and non-permissive energetic minimum.

To test this hypothesis and resolve the structural discrepancies between experimental and AI-predicted GLUT4 models, we implemented a comprehensive computational pipeline linking equilibrium simulations with non-equilibrium steered molecular dynamics (SMD). This strategy allowed us to quantify non-equilibrium work accumulation (*W*_raw_) and reconstruct the underlying free-energy landscapes of hexose translocation. We systematically evaluated the apo, ADP-bound, and ATP-bound states across physiological (310.15 K) and reduced (303.15 K) temperatures to isolate the direct interplay between pore geometry, backbone stability, and steric desolvation penalties. To expose the underlying allosteric pathways, we mapped internal signal propagation using ensemble cross-correlation heatmaps and graph-theoretical network analysis. Crucially, this integrated biophysical framework provides a rigorous diagnostic of the dynamic fidelity inherent to native Cryo-EM templates relative to AlphaFold predictions. We demonstrate that while native structures preserve the conformational plasticity and dynamic hydration shells needed for efficient glucose transport, the over-optimized packing of predictive models imposes severe mechanical and kinematic gridlock.

## 2. Results

### 2.1. Ligand Interactions

Allosteric ligands modulate membrane transport proteins by inducing conformational changes that stabilize or destabilize specific states, unlike inhibitors that directly block transport. Characterizing the effects of ligands like ATP and ADP is vital for GLUT4, as their interactions reshape the free energy barriers of glucose translocation, illuminating how intracellular biochemical conditions govern transport behaviour.

#### 2.1.1. Adenosine Tri-Phosphate (ATP) Interactions

Comparative structural analysis reveals both conserved purine recognition motifs and prominent topological divergences within the ATP-binding region of GLUT4 (Figure 3). Adenine base coordination is broadly preserved across both structural models via hydrogen-bonding matrices anchored at the roof of the binding pocket, mediated by **THR351** and **GLY348** in the AlphaFold template and **ARG454** and **GLY453** in the experimental Cryo-EM structure. However, significant divergence occurs surrounding the ribose core. The native Cryo-EM structure forms a dense coordination network around the ribose moiety, stabilizing the nucleoside through multivalent hydrogen bonds and electrostatic contacts with **ARG329**, **ARG330**, and **THR331**. In contrast, the AlphaFold model leaves the ribose ring substantially solvent-exposed, reflecting unoptimized side-chain packing and incomplete local structural complementarity.

**Figure 3.**
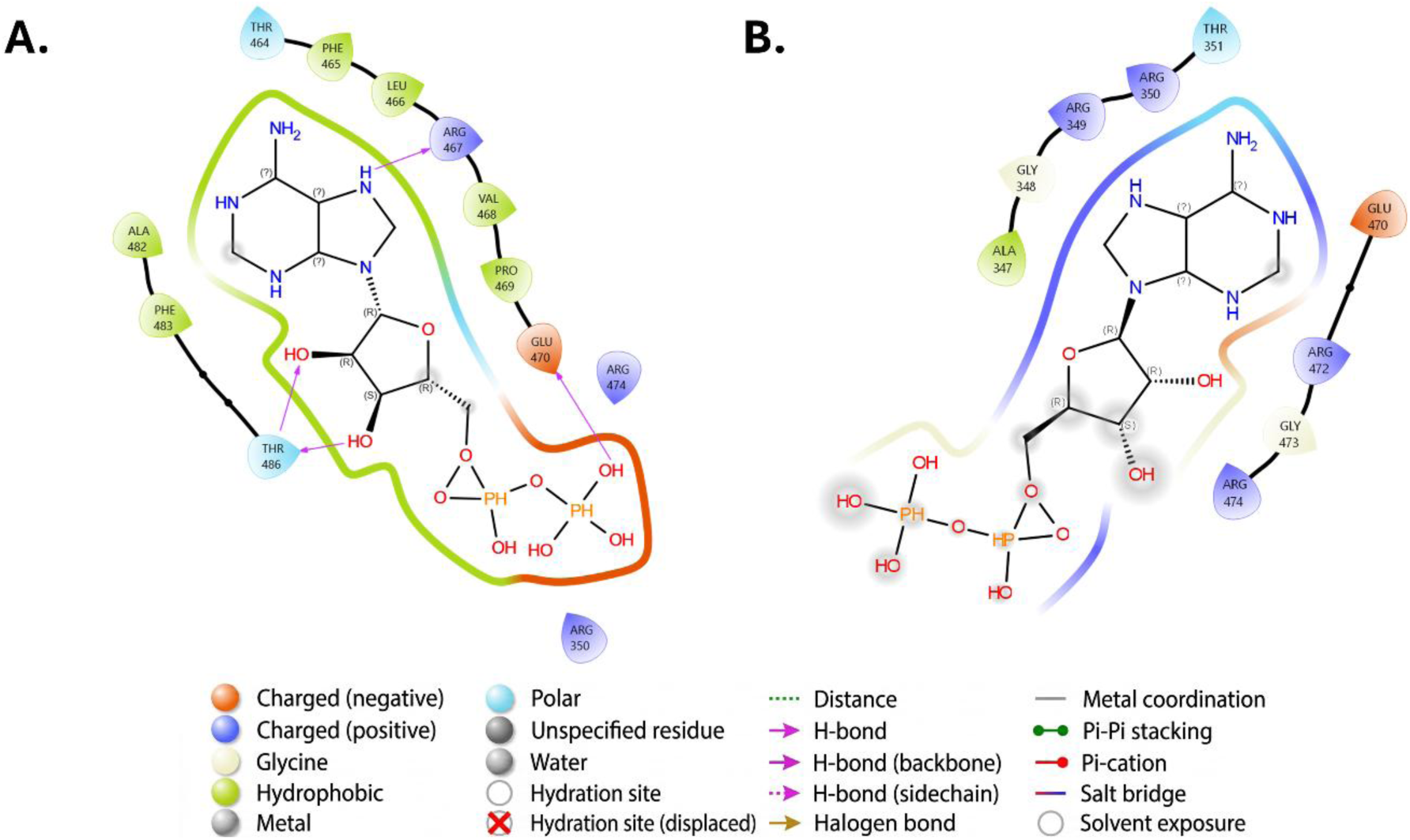
Interaction diagram for ATP, **A** : AlphaFold model, **B** : CryoEM model.

Further discrepancies emerge across the coordination environment of the highly electronegative triphosphate tail. While the AlphaFold predicted model stabilizes the phosphate group primarily through discrete electrostatic pairings with **ARG472** and **ARG474**, the experimental Cryo-EM framework constructs an extensive coordination nest involving **ARG447**, **ARG462**, and **PHE445**. This multivalent coordination nest generates an intricate network of electrostatic and steric contacts that neutralizes negative charge density while facilitating induced-fit side-chain rearrangements. Consequently, the native Cryo-EM structure captures an adaptive binding architecture optimized for ligand accommodation, whereas the static AlphaFold template maintains an apo-like binding cavity that lacks these dynamic conformational adjustments.

#### 2.1.2. Adenosine Di-Phosphate (ADP) Interactions

Adenosine diphosphate (ADP) lacks the terminal *γ*-phosphate, reducing its net physiological charge to −2 and substantially lowering the total electrostatic demand on the orthosteric binding pocket (Figure 4). While hydrophobic and hydrogen-bonding contacts for adenine base recognition remain conserved between both models, coordination of the ribose core displays a topological reversal relative to ATP. Within the AlphaFold predicted template, the ribose hydroxyl groups undergo direct, bivalent hydrogen bonding with the side chain of **THR486**. Conversely, the native Cryo-EM pocket leaves the ribose sugar largely solvent-exposed, reflecting local conformational relaxation and side-chain uncoupling from the nucleoside core following the loss of the terminal phosphate.

**Figure 4.**
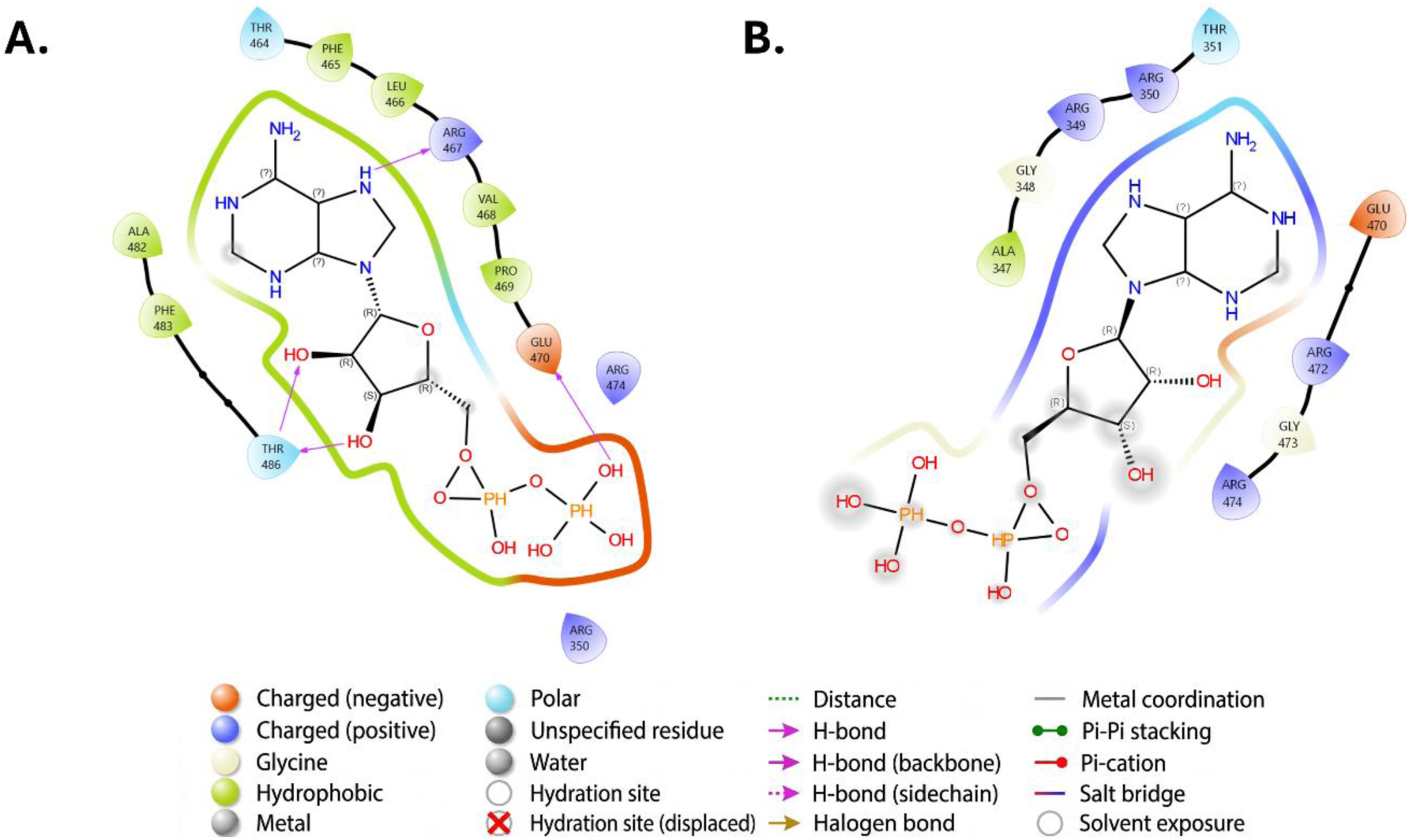
Interaction diagram for ATP, **A** : AlphaFold model, **B** : CryoEM model

The coordination profile of the truncated diphosphate tail highlights distinct charge-stabilization strategies. The AlphaFold model stabilizes the terminal phosphate groups through targeted electrostatic interactions involving **ARG474** and **GLU470** while maintaining elevated solvent accessibility at the pocket periphery. In contrast, the experimental Cryo-EM structure coordinates the diphosphate moiety through a localized polar network comprising **ARG472**, **ARG474**, and **GLY473**. Notably, this experimental pocket completely lacks the dense hydrophobic scaffold that cradled the larger triphosphate tail of ATP, confirming a local structural reset upon ADP binding. Because ADP imposes less total charge density and conformational strain on the cytoplasmic domain, the static AlphaFold model serves as a closer approximation of the native ADP-bound state than it does for the ATP-bound complex.

**Figure 5.**
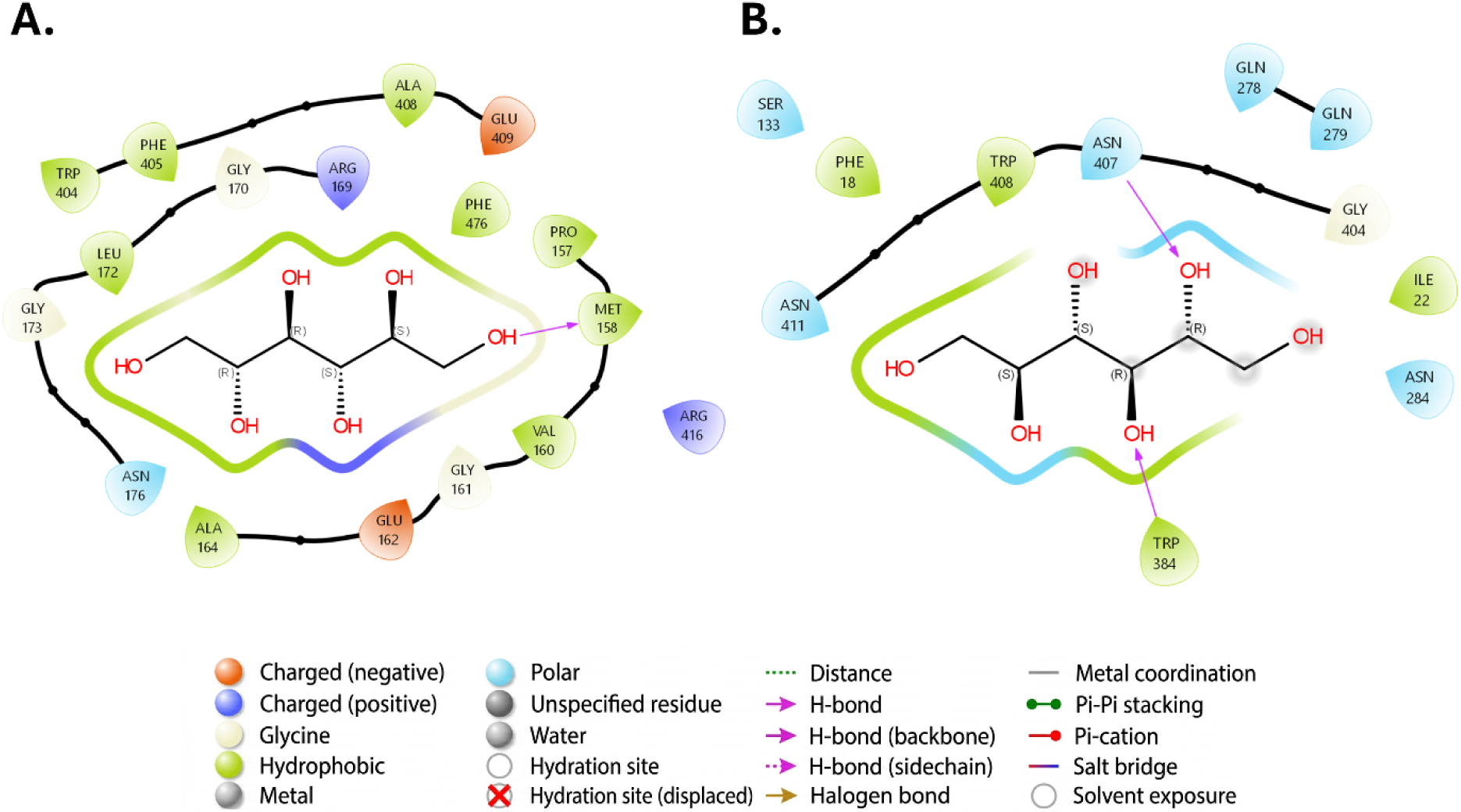
Interaction diagram for DEX, **A** : AlphaFold model, **B** : CryoEM model

#### 2.1.3. Dextrose (DEX) Interactions

As a neutral six-carbon polyol, glucose recognition within GLUT4 relies on directional hydrogen bonding and transient nonpolar interactions rather than long-range electrostatics. Comparative analysis reveals critical topological discrepancies between the predicted AlphaFold model and the experimental Cryo-EM structure.

Within the AlphaFold template, the pyranose ring is enclosed in a dense hydrophobic shell (**TRP404**, **PHE405**, **LEU172**, **MET158**), reflecting an inwardly collapsed, apo-like cavity optimized for intramolecular packing without solvent stabilization. In stark contrast, the native Cryo-EM structure features a more open spatial arrangement (**TRP408**, **PHE18**), establishing a solvent-accessible, transport-competent environment engineered for rapid substrate entry and transient stabilization.

Polar recognition further differentiates the structural models. Hydrogen bonding in AlphaFold is structurally isolated, relying on localized interactions with **ARG169** and backbone contacts with **MET158**. Conversely, the native Cryo-EM framework organizes an extensive, high-fidelity polar lattice comprising **ASN407**, **ASN411**, **GLN278**, **GLN279**, and **TRP384**. This distributed network provides precise stereochemical complementarity to the hydroxyl-rich surface of hexose, enabling dynamic, low-barrier interactions essential for facilitated diffusion.

Collectively, these localized orthosteric interaction profiles establish the initial energetic boundary conditions and physical constraints governing global protein stability, thermal fluctuations, and non-equilibrium transport mechanics across the transporter matrix.

### 2.2. Conformational Dynamics

#### 2.2.1. Apo State

**Figure 6.**
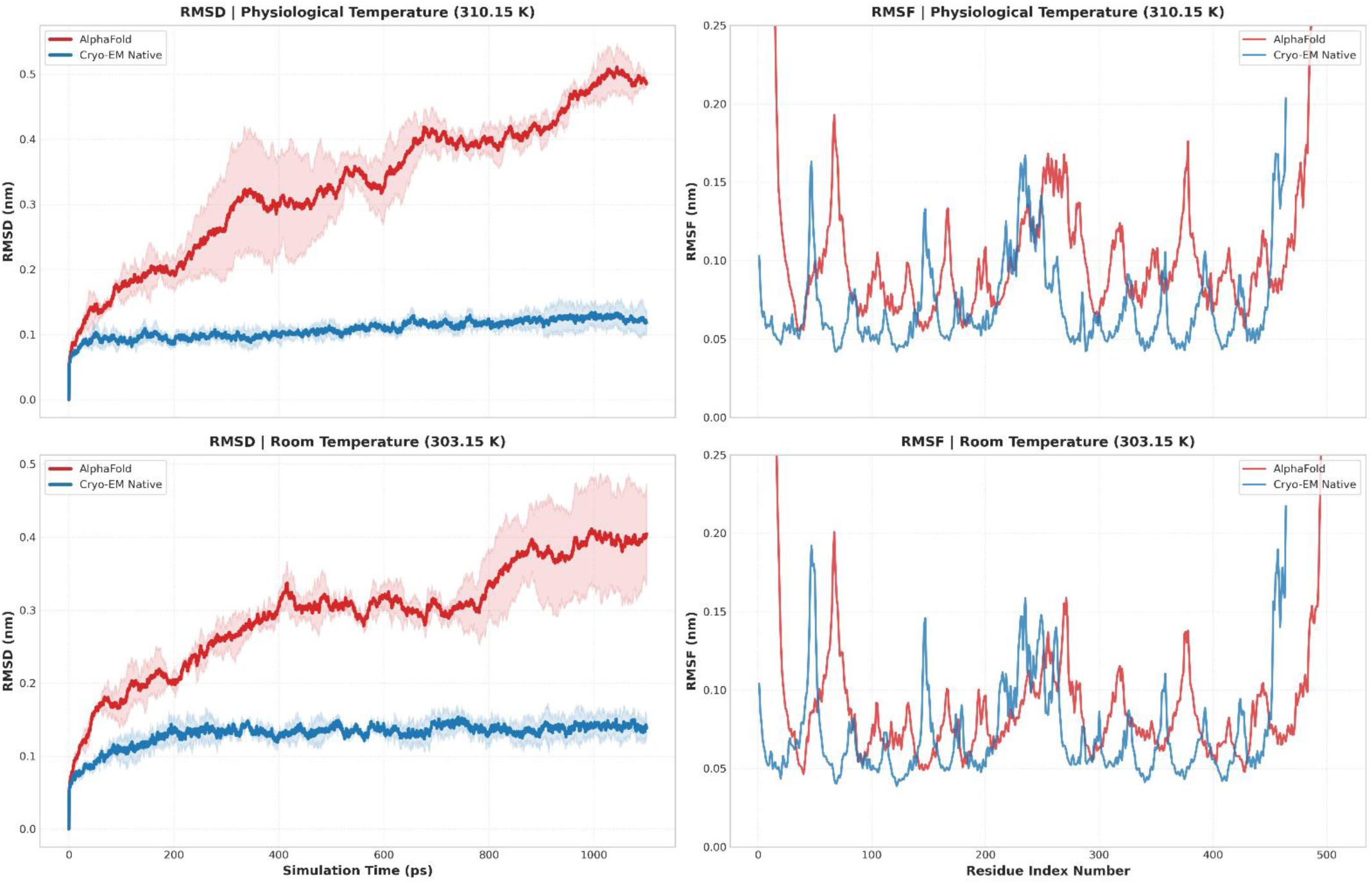
Conformational Stability of Apo GLUT4. Backbone root-mean-square deviation (RMSD; left) and residue-wise root-mean-square fluctuation (RMSF; right) of the Cryo-EM (blue) and AlphaFold (red) models under physiological (310.15 K; top) and reduced (303.15 K; bottom) temperatures. Solid lines represent the mean trajectories, while shaded regions indicate the standard deviation across independent simulations.

Under physiological boundary conditions (**310.15 K**), the unliganded Cryo-EM structure exhibits high global backbone stability, equilibrating within **50 ps** and maintaining a flat backbone Root Mean Square Deviation (RMSD) plateau between **0.10 nm** and **0.13 nm** across the entire **2 ns** trajectory. Conversely, the AlphaFold template displays continuous structural drift, breaching **0.20 nm** by **150 ps** and escalating steadily to a terminal maximum of **0.50 ± 0.03 nm** with broad replica variance. Thermal attenuation to **303.15 K** curtails AlphaFold’s terminal drift to **0.40 ± 0.07 nm** (following a metastable plateau from **500 ps** to **1500 ps**), whereas the Cryo-EM framework tracks flat at ∼**0.13 nm**. This persistent divergence confirms that the global instability of the predicted model stems from an intrinsic topological defect rather than stochastic thermal fluctuations. Residue-wise Root Mean Square Fluctuation (RMSF) mapping isolates the atomistic coordinates driving this global divergence. Solvent-exposed N- and C-terminal loops in the AlphaFold structure undergo large-amplitude movements exceeding **0.25 nm** across both temperatures, whereas native Cryo-EM terminal extremities remain tightly bounded below **0.12 nm**. While the extracellular loop domain (residues 50–70) acts as a shared flexible hotspot in both models (**0.16–0.19 nm**), a critical mechanical divergence occurs within the primary translocation core (residues 230–280). The native Cryo-EM backbone maintains a cohesive, low-amplitude baseline of **0.05–0.08 nm**, preserving localized channel patency. In contrast, the AlphaFold template exhibits severe local instability throughout this central cavity, generating broad fluctuation peaks that reach **0.15–0.17 nm** across both thermal regimes.

These unliganded dynamics confirm that the native Cryo-EM framework preserves structural integrity and spatial patency across the unconstrained transport pathway. Conversely, the high core volatility and continuous backbone drift in the predicted AlphaFold template reflect an intrinsically unstable, collapsed apo state incapable of supporting low-barrier alternating-access transport mechanics.

#### 2.2.2. ATP-Bound State

**Figure 7.**
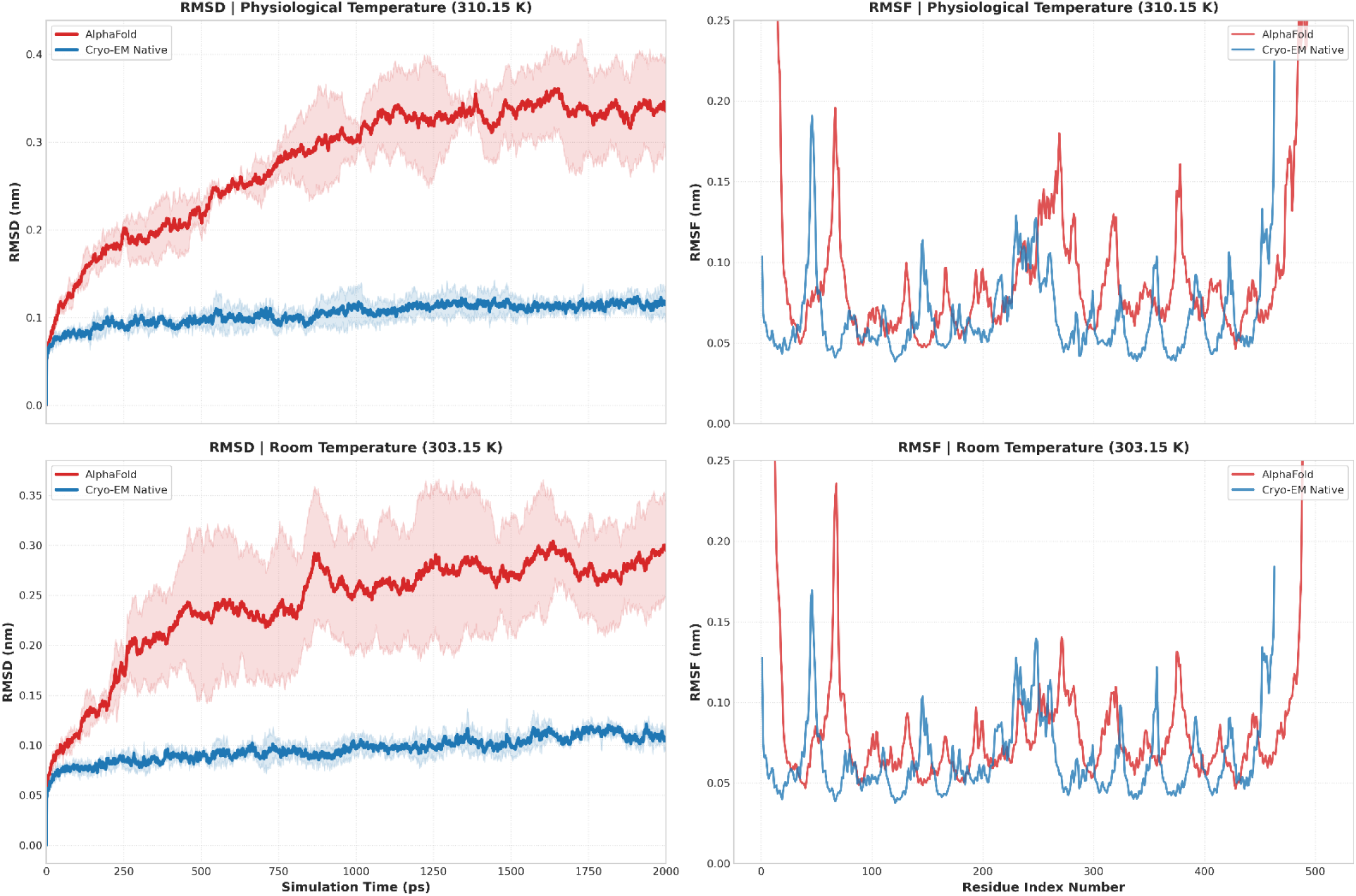
Conformational Stability of ATP-Bound GLUT4. Backbone root-mean-square deviation (RMSD; left) and residue-wise root-mean-square fluctuation (RMSF; right) of the Cryo-EM (blue) and AlphaFold (red) models in the ATP-bound state under physiological (310.15 K; top) and reduced (303.15 K; bottom) temperatures. Solid lines represent the mean trajectories, while shaded regions indicate the standard deviation across independent simulations.

Under physiological conditions (**310.15 K**), the ATP-bound Cryo-EM native backbone maintains excellent global stability, rapidly equilibrating within **50 ps** to establish an asymptotic RMSD plateau strictly bounded between **0.08 nm** and **0.12 nm** over **2 ns**. In stark contrast, the AlphaFold template undergoes continuous kinematic drift under active ligand modulation, breaching **0.20 nm** by **350 ps** and terminating at a maximum plateau of **0.33–0.35 nm** with substantial ensemble variance. Upon thermal attenuation to **303.15 K**, the Cryo-EM structure displays minimal thermal sensitivity, tracking along the same **0.08–0.12 nm** baseline. Although thermal cooling curtails AlphaFold’s terminal drift to **0.28–0.30 nm** (following a volatile metastable phase between **0.23 nm** and **0.28 nm**), it fails to globally stabilize the predicted template.

Residue flexibility mapping highlights profound differences in local fold containment during ATP binding. Solvent-exposed N- and C-terminal extremities in the AlphaFold template spike aggressively out of the viewing window (**>0.25 nm**), whereas native Cryo-EM loops remain anchored below **0.12 nm**. Both models capture extracellular loop flexibility (residues 50–70) peaking between **0.17 nm** and **0.23 nm**. However, deep within the primary translocation core (residues 230–280), the native Cryo-EM backbone preserves an exceptionally rigid configuration oscillating tightly between **0.04 nm** and **0.08 nm** to maintain channel patency under nucleotide stress. Conversely, the AlphaFold core displays widespread kinematic disruption, with disorganized local fluctuation peaks spiking to **0.15–0.18 nm** at **310.15 K** and persisting at **0.12–0.14 nm** at **303.15 K**.

Multivalent ATP coordination in native GLUT4 stabilizes the core gating domain to maintain a transport-competent conformation under nucleotide stress. The widespread core disruption and persistent global drift in AlphaFold demonstrate that the computational model fails to translate ATP binding into productive allosteric stabilization, resulting in structural uncoupling.

#### 2.2.3. ADP-Bound State

**Figure 8.**
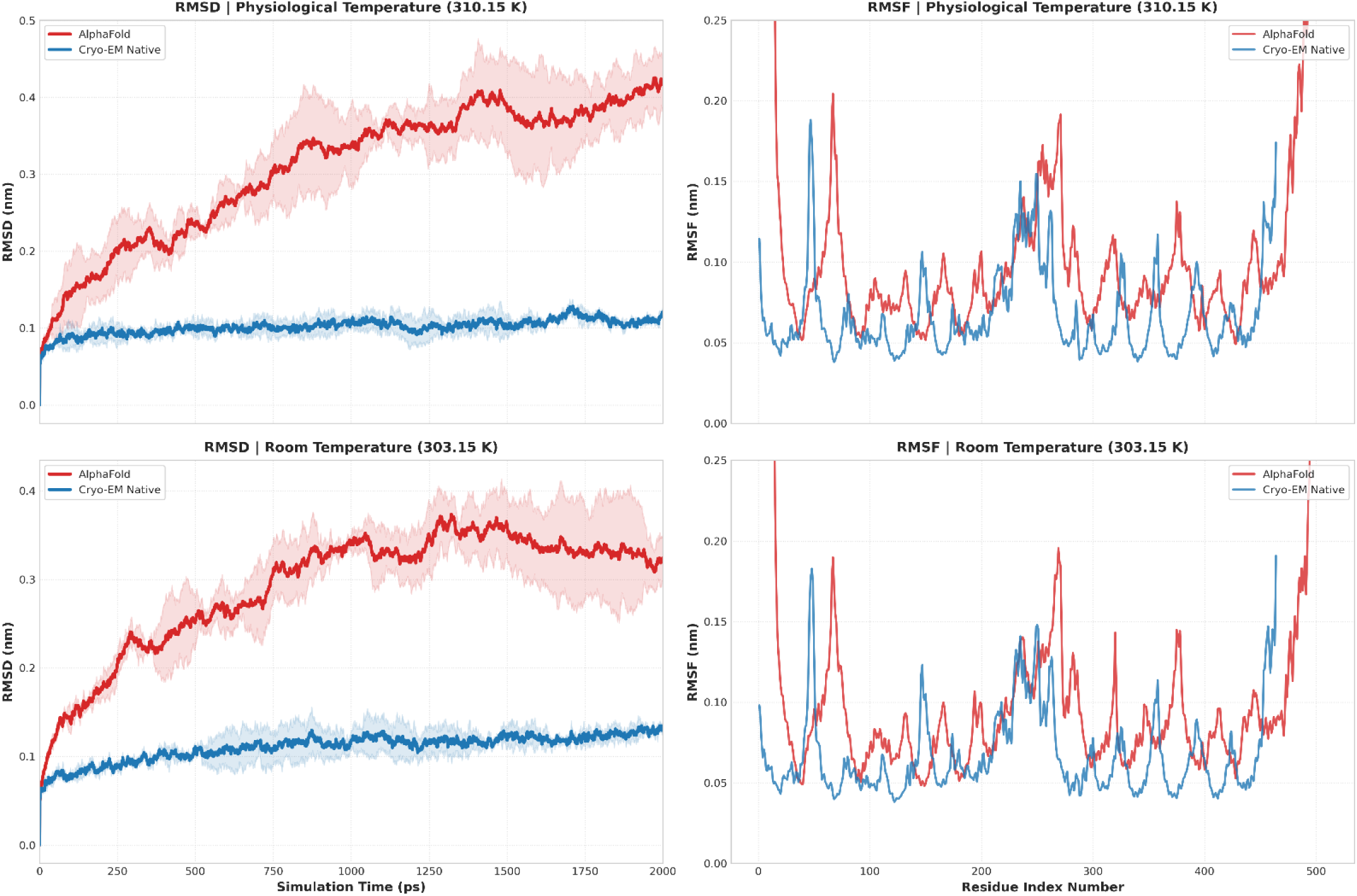
Conformational Stability of ADP-Bound GLUT4. Backbone root-mean-square deviation (RMSD; left) and residue-wise root-mean-square fluctuation (RMSF; right) of the Cryo-EM (blue) and AlphaFold (red) models in the ADP-bound state under physiological (310.15 K; top) and reduced (303.15 K; bottom) temperatures. Solid lines represent the mean trajectories, while shaded regions indicate the standard deviation across independent simulations.

At a physiological temperature of **310.15 K**, the ADP-bound Cryo-EM framework demonstrates robust global stability, equilibrating within **50 ps** to hold a tightly bounded RMSD plateau between **0.08 nm** and **0.12 nm** across **2 ns**. Conversely, the AlphaFold model displays continuous, non-asymptotic structural drift, breaching the **0.20 nm** threshold by **400 ps** and climbing to a terminal maximum of **0.40–0.43 nm** with broad ensemble variance. Under room temperature conditions (**303.15 K**), the experimental structure preserves its low-amplitude baseline (**0.08–0.14 nm**), proving minimal thermal sensitivity. The AlphaFold model drifts to **0.25 nm** by **500 ps**, climbs to a metastable peak of **0.35–0.37 nm** at **1400 ps**, and terminates at **0.32–0.35 nm**, demonstrating persistent macroscale instability under ADP modulation.

Local flexibility profiles confirm a stark divergence in backbone cohesion during ADP coordination. Solvent-accessible N- and C-terminal loops in the AlphaFold structure exhibit unconstrained movements exceeding **0.25 nm** across both thermal regimes, whereas native Cryo-EM extremities remain tightly bounded below **0.12 nm**. Extracellular loop fluctuations (residues 50–70) align closely between both models at **0.18–0.21 nm**. Crucially, within the central translocation core (residues 230–280), the native Cryo-EM backbone preserves a rigid baseline oscillating between **0.04 nm** and **0.09 nm** to maintain core channel patency. Conversely, the AlphaFold template demonstrates severe kinematic disruption across this core sequence, generating broad, multi-peaked fluctuations that reach **0.16–0.19 nm** across both temperature regimes.

These dynamics demonstrate that native ADP binding maintains a stable, low-strain translocation core identical to the unliganded state, supporting efficient substrate exchange upon loss of the terminal *γ*-phosphate. In contrast, the AlphaFold template suffers systematic core breakdown, misrepresenting the conformational relaxation and dynamic patency of the native ADP-bound state.

### 2.3. Work profiles of Glucose Transport

To evaluate the functional energetic barriers governing substrate translocation, non-equilibrium steered molecular dynamics (SMD) simulations quantified the cumulative mechanical work (*W*_raw_) required to drive glucose through the central pore over a 2.0 nm pulling coordinate. Comparing time-resolved work profiles across Apo, ATP-bound, and ADP-bound ensembles under physiological (310.15 K) and reduced (303.15 K) temperatures isolates the steric and thermodynamic constraints imposed by nucleotide coordination, providing a direct physical measure of channel friction and translocation kinetics.

#### 2.3.1. ATP-Bound State

**Figure 9.**
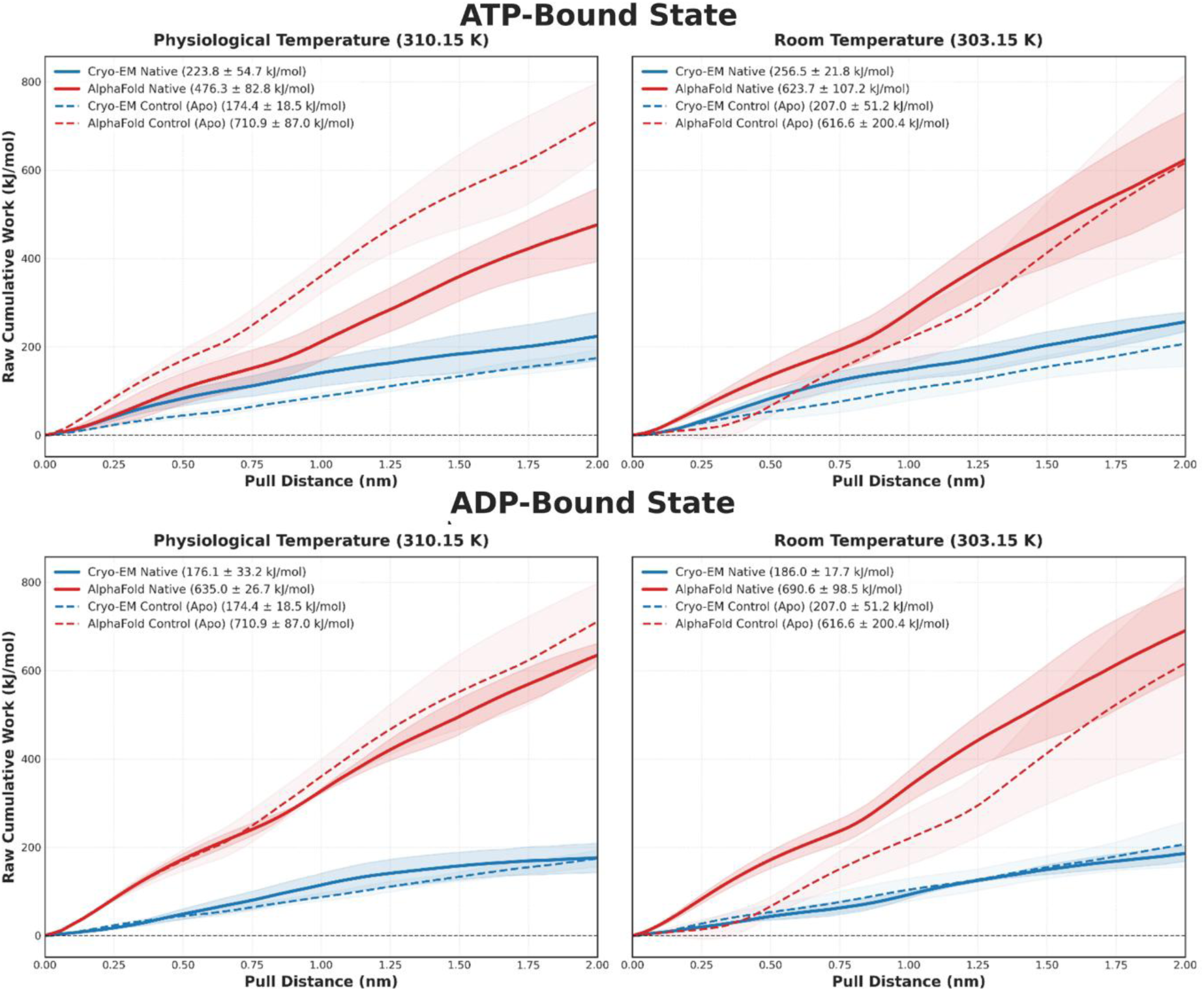
Work Profiles of Glucose Translocation in the ATP & ADP-Bound State. Cumulative work profiles (Wraw) for glucose translocation through the Cryo-EM (blue) and AlphaFold (red) models under physiological (310.15 K; left) and reduced (303.15 K; right) temperatures. Solid lines represent ATP-bound simulations, whereas dashed lines indicate the corresponding apo control simulations. Shaded regions represent the standard deviation across independent triplicate simulations. Final cumulative work values are reported in the legend.

Under physiological conditions (310.15 K), the experimentally resolved Cryo-EM framework operates within a low-energy, transport-competent regime. The unliganded Apo control requires a minimal work expenditure of 174.4 ± 18.5 kJ/mol, while ATP coordination induces a modest, controlled increment to 223.8 ± 54.7 kJ/mol. Thermal attenuation to 303.15 K causes a predictable, uniform scale-up in mechanical resistance to 207.0 ± 51.2 kJ/mol (Apo) and 256.5 ± 21.8 kJ/mol (ATP), maintaining tightly bounded variance envelopes across the entire 2.0 nm axis.

Conversely, the predicted AlphaFold model imposes extreme thermodynamic resistance. At 310.15 K, the unliganded Apo control reaches a steep 710.9 ± 87.0 kJ/mol, reflecting severe steric gridlock within the collapsed pore. ATP binding partially lowers this barrier to 476.3 ± 82.8 kJ/mol. At 303.15 K, however, AlphaFold exhibits severe thermal volatility and erratic trajectory inversions, escalating to 623.7 ± 107.2 kJ/mol (ATP) and 616.6 ± 200.4 kJ/mol (Apo) alongside a massive standard deviation envelope.

Comparing these profiles reveals that while native ATP binding executes a low-barrier, elastic clamping event (223.8 kJ/mol at 310.15 K) that preserves an open conduction pathway, the AlphaFold template subjects the substrate to a massive energy penalty (> 470 kJ/mol) driven by physical pore constriction and uncoordinated structural collapse.

#### 2.3.2. ADP-Bound State

In the ADP-bound state at 310.15 K, the native Cryo-EM framework maintains an exceptionally low, uniform energetic landscape. Tracking almost identically to its corresponding Apo control (174.4 ± 18.5 kJ/mol), the ADP complex culminates in a terminal work value of 176.1 ± 33.2 kJ/mol. Upon thermal reduction to 303.15 K, work expenditures scale smoothly to 186.0 ± 17.7 kJ/mol (ADP) and 207.0 ± 51.2 kJ/mol (Apo), preserving overlapping variance envelopes throughout the pulling trajectory.

In contrast, the ADP-bound AlphaFold model perpetuates severe mechanical resistance. At 310.15 K, ADP coordination marginally reduces the extreme Apo work barrier (710.9 ± 87.0 kJ/mol) to 635.0 ± 26.7 kJ/mol, leaving the pore locked in a high-resistance state. Thermal attenuation to 303.15 K worsens this mechanical demand, driving the ADP trajectory to 690.6 ± 98.5 kJ/mol and the Apo control to 616.6 ± 200.4 kJ/mol alongside substantial trajectory divergence.

While native ADP coordination preserves an unhindered, low-friction pathway matching the unliganded state (176.1 kJ/mol vs 174.4 kJ/mol), AlphaFold remains locked in a high-resistance conformation (> 630 kJ/mol). This confirms that loss of the terminal *γ*-phosphate in native GLUT4 restores open channel patency, whereas the computational fold fails to model this dynamic conformational relaxation.

### 2.4. Interaction Energies

To delineate the energetic determinants governing hexose translocation, non-bonded interaction potentials between glucose and the GLUT4 matrix were decomposed into short-range electrostatic (Coulombic) and van der Waals (Lennard-Jones) potentials during steered molecular dynamics (SMD). This decomposition resolves how nucleotide occupancy shapes local chemical forces, mapping the steric and polar constraints that drive macroscale work variations.

#### 2.4.1. Apo State

**Figure 10.**
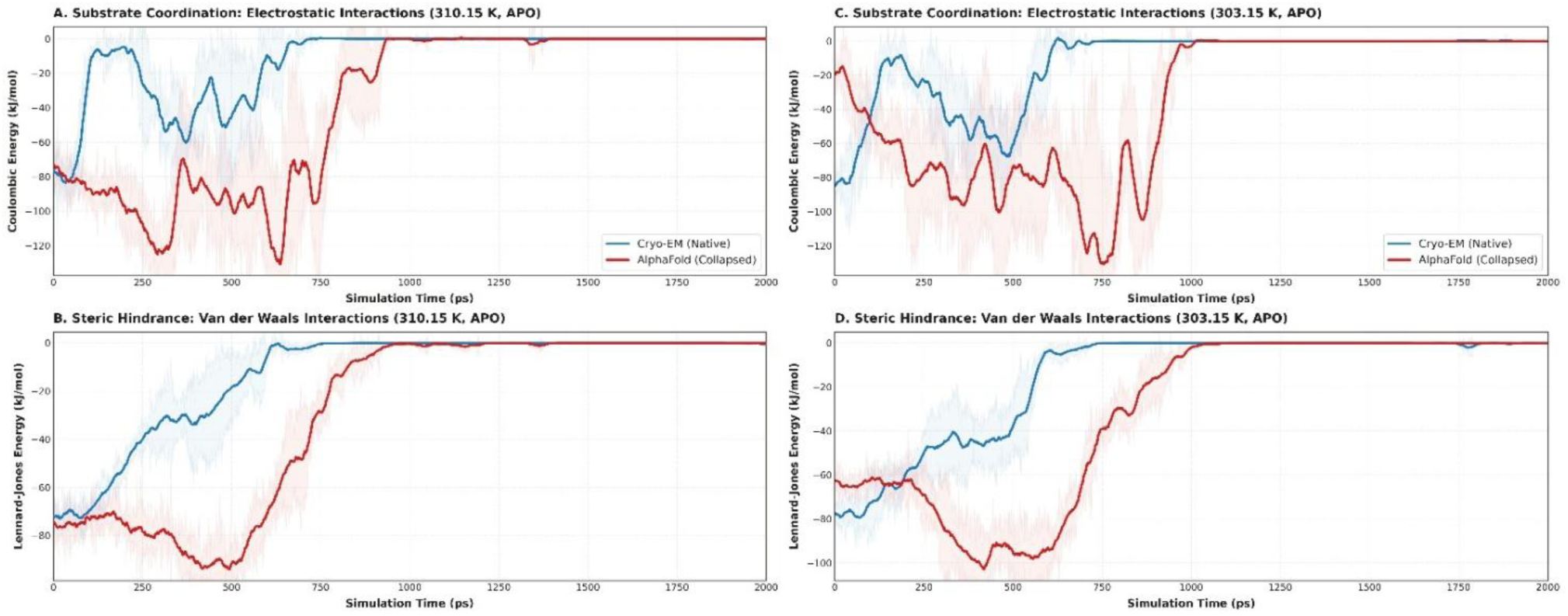
Interaction Energy Profiles of Apo GLUT4. Electrostatic (A) and van der Waals (B) interaction energies between glucose and GLUT4 during steered molecular dynamics simulations of the apo state under physiological (310.15 K; left) and reduced (303.15 K; right) temperatures. The Cryo-EM model is shown in blue and the AlphaFold model in red. Solid lines represent the mean interaction energies, while shaded regions indicate the standard deviation across independent triplicate simulations.

At **310.15 K**, electrostatic energy profiles reveal an efficient, low-barrier polar path in the native Cryo-EM structure. An initial potential of **-75 to -80 kJ/mol** rapidly relaxes to **-10 kJ/mol** within **100 ps**, with minor transient stabilization events (**-50 kJ/mol**) clearing completely by **800 ps**. Under thermal attenuation (**303.15 K**), the native framework preserves this low-resistance baseline, clearing by **650 ps**. In contrast, the AlphaFold template subjects the hexose to severe polar entrapment across both temperatures. Initiating at **-85 kJ/mol**, the predicted trajectory plunges into deep electrostatic valleys reaching global minima of **-120 to -130 kJ/mol** (**320–750 ps**), delaying baseline clearance until **950–1000 ps**.

Lennard-Jones interaction timelines at **310.15 K** isolate the steric friction driving translocation. The native Cryo-EM framework exhibits a smooth, monotonic decay of non-bonded contact forces, ascending continuously from **-72 kJ/mol** to complete baseline dissipation by **750 ps**, quantifying an unobstructed passage through a patulous pore. At **303.15 K**, it scales similarly to reach baseline by **700 ps**. Conversely, the AlphaFold model records a severe, deterministic steric bottleneck. Trajectories at both **310.15 K** and **303.15 K** plunge into a prolonged confinement valley holding a maximum contact trap of **-95 to -100 kJ/mol** (**400–500 ps**), preventing baseline clearance until **1000 ps**.

#### 2.4.2. ATP-Bound State

**Figure 12.**
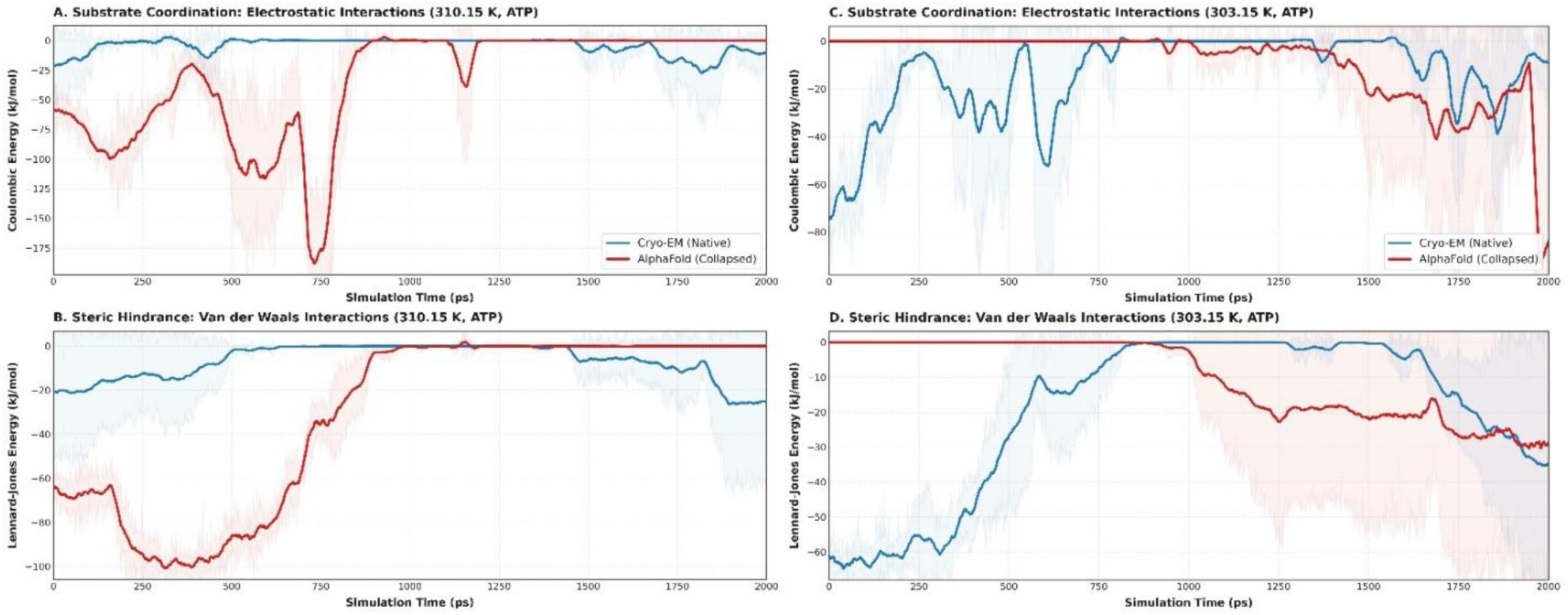
Interaction Energy Profiles of ATP-Bound GLUT4.

During physiological ATP occupancy (**310.15 K**), the native Cryo-EM template displays early, low-amplitude polar coordination (**-20 kJ/mol**) that relaxes to zero by **500 ps**. Under reduced temperature (**303.15 K**), it undergoes minor, bounded fluctuations (**-5 to -40 kJ/mol**) before clearing cleanly at **650 ps**. Conversely, the AlphaFold template causes extreme electrostatic entrapment at **310.15 K**, plunging into an asymmetric double-well profile with a catastrophic global minimum of **-175 kJ/mol** at **720 ps**. At **303.15 K**, AlphaFold displays an anomalous **1.35 ns** interaction lag phase (**0 kJ/mol**), followed by late-stage chaotic plunges reaching **-90 kJ/mol**.

Short-range Lennard-Jones profiles at **310.15 K** show that native Cryo-EM maintains a low-friction pathway (**-20 kJ/mol** initial), ascending to the zero-interaction baseline by **550 ps**. At **303.15 K**, van der Waals contacts persist slightly longer (**-60 kJ/mol** initial) before clearing at **850 ps**. In sharp contrast, the AlphaFold template at **310.15 K** drops into a deep confinement well holding a maximum contact plateau of **-100 kJ/mol** (**250–450 ps**). At **303.15 K**, it replicates its electrostatic lag behavior (**0 kJ/mol** for **1000 ps**) before stepping down to late-stage contact values of **-30 kJ/mol**.

#### 2.4.3. ADP Bound State

**Figure 13.**
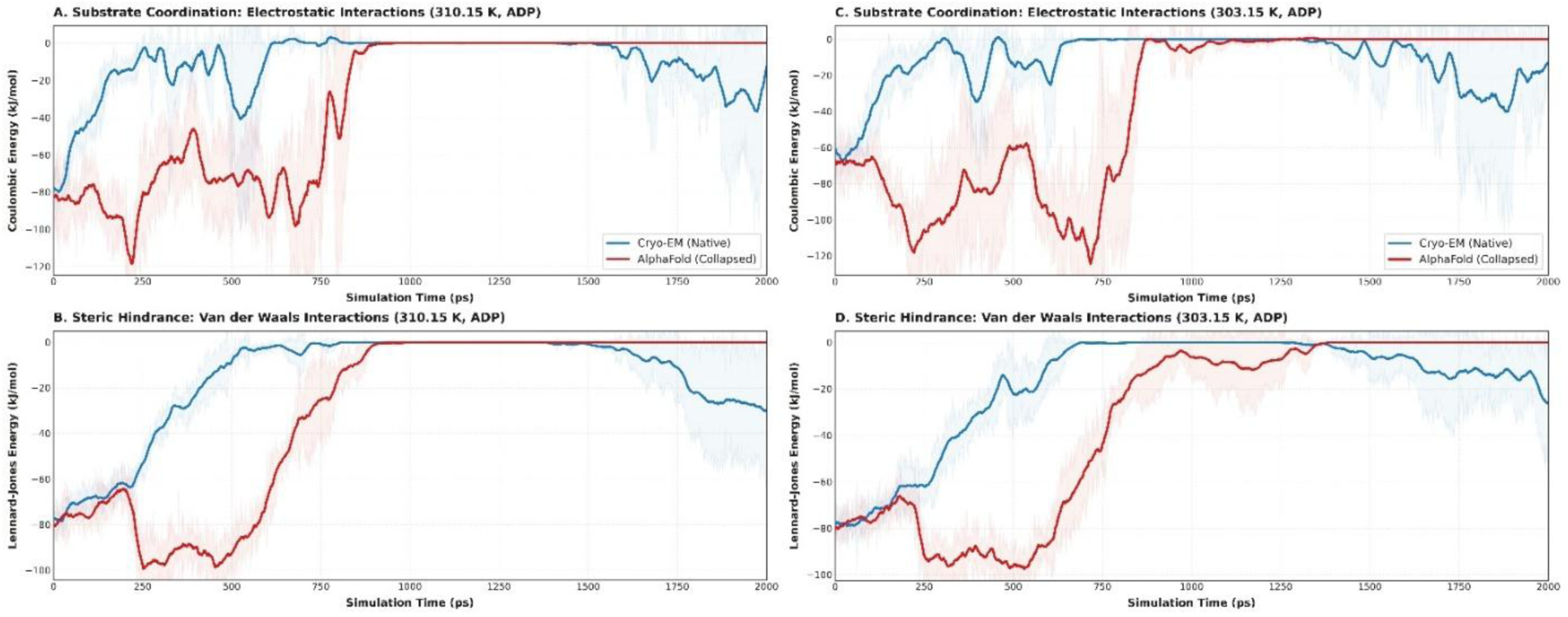
Interaction Energy Profiles of ADP-Bound GLUT4.

At **310.15 K**, the ADP-bound Cryo-EM framework establishes a low-barrier polar path, rising from **-60 kJ/mol** to **-15 kJ/mol** within **200 ps** and clearing completely by **700 ps**. At **303.15 K**, it initiates at **-90 kJ/mol**, climbs swiftly to **-20 kJ/mol**, and holds a clean baseline until minor late-stage re-coordination. Conversely, the AlphaFold model demonstrates intense electrostatic entrapment across both temperatures. Initiating at **-60 to -70 kJ/mol**, trajectories plunge into severe local minima of **-115 to -120 kJ/mol** (**250–720 ps**), delaying baseline clearance to **850 ps**.

Lennard-Jones profiles at **310.15 K** show native Cryo-EM ascending smoothly from **-80 kJ/mol** to zero baseline by **600 ps**, holding flat thereafter. At **303.15 K**, it ascends progressively to reach baseline by **700 ps**. In contrast, the AlphaFold model documents an immediate, temperature-invariant steric trap. Trajectories at both **310.15 K** and **303.15 K** plunge into an identical, extended confinement valley holding a maximum contact plateau of **-95 to -100 kJ/mol** (**250–550 ps**) before ascending to baseline at **850 ps**.

### 2.5. Pore geometry Analysis

Spatial pore radius profiles computed using the Channel Annotation Package (CHAP) quantify the structural patency, bottleneck dimensions, and gating shifts of the translocation pathway across Apo, ATP-bound, and ADP-bound ensembles under physiological (310.15 K) and thermally attenuated (303.15 K) boundary conditions.

**Figure 17.**
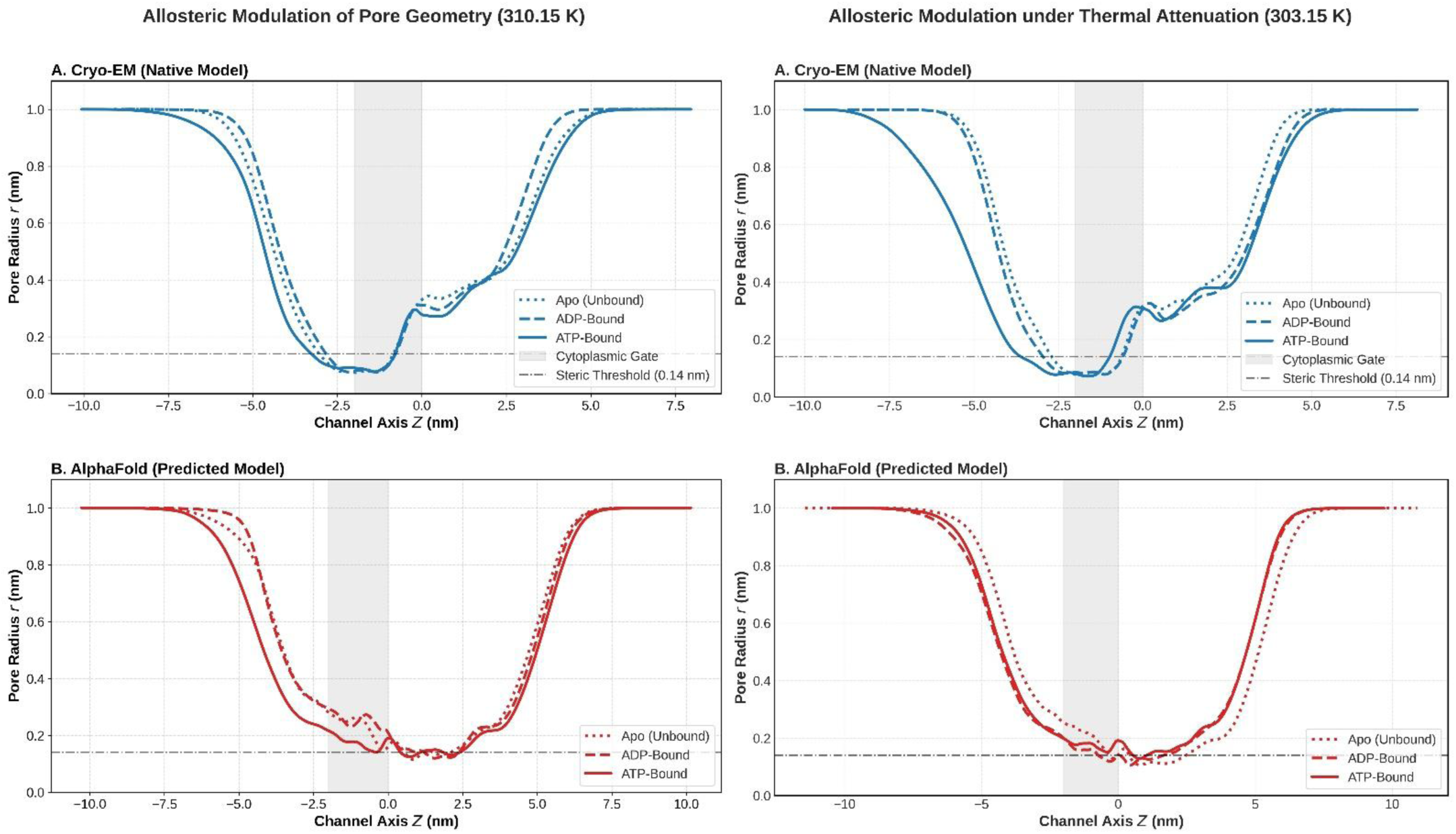
Pore Geometry of GLUT4 under Physiological Conditions (310.15 K). Pore radius profiles calculated using the Channel Annotation Package (CHAP) along the principal channel axis for the Cryo-EM (A) and AlphaFold (B) models under physiological temperature (310.15 K). Superimposed profiles of the apo (dotted), ADP-bound (dashed), and ATP-bound (solid) states illustrate ligand-dependent modulation of the transport pathway. The shaded region denotes the cytoplasmic gate, while the horizontal dash–dot line indicates the steric threshold (0.14 nm) for pore hydration.

In the experimentally resolved structure, the central permeation pathway maintains an open, transport-competent extracellular entry and luminal vestibule before transitioning into a sharply localized constriction at the designated cytoplasmic gate. Across all functional states, the pore radius plunges decisively below the 0.14 nm hydration threshold, establishing a tight bottleneck radius minimum of 0.05 nm. Gate shifts along the intracellular exit flank reveal clear allosteric modulation driven by nucleotide occupancy: ADP stabilizes the widest exiting aperture (0.38 nm), Apo maintains an intermediate opening (0.28 nm), and ATP enforces the tightest intracellular funneling (0.18 nm).

The computational model exhibits a severe topological dislocation of its primary gating mechanism. Rather than constricting at the cytoplasmic gate, the primary physical bottleneck shifts extracellularly into the central channel lumen, where the pore radius drops to 0.10–0.14 nm—breaching the hydration threshold in a region where native GLUT4 remains widely open (0.35–0.50 nm). Furthermore, the predicted cytoplasmic gate fails to execute uniform closure: ADP remains abnormally dilated (0.30 nm), Apo fluctuates loosely (0.18–0.32 nm), and ATP forms only an ill-defined secondary constriction (0.12–0.25 nm).

Thermal attenuation preserves the localized cytoplasmic gating architecture of the native framework while enhancing allosteric differentiation. The luminal vestibule remains widely open (0.30–0.48 nm) before the profile plunges sharply at the cytoplasmic gate to hit an identical bottleneck radius minimum of 0.05 nm across all states. Gate shift dynamics along the intracellular exit flank reveal that ATP opens significantly earlier (expanding to 0.38 nm), whereas Apo and ADP remain tightly constricted prior to terminal expansion.

Under reduced thermal energy, the predicted model perpetuates a distorted, non-functional pore geometry. The primary physical bottleneck remains abnormally shifted into the central channel lumen, breaching the 0.14 nm hydration threshold down to 0.10–0.14 nm. At the true cytoplasmic gate, the computational fold fails to establish structural closure: Apo stays overly open (0.30 nm), while ATP and ADP display shallow, featureless radius fluctuations (0.12–0.25 nm) lacking distinct state-dependent gate shifts.

### 2.6. Solvent Accessibility and Hydration Shell Dynamics

Hexose translocation requires dynamic aqueous lubrication through the channel lumen. Tracking the primary hydration shell of glucose (*N*_water_ ≤ 0.5 nm) across SMD trajectories (Cryo-EM vs. AlphaFold; Apo/ATP/ADP; 310.15/303.15 K) resolves whether transport barriers stem from functional desolvation or collapse-induced dewetting.

**Figure 19.**
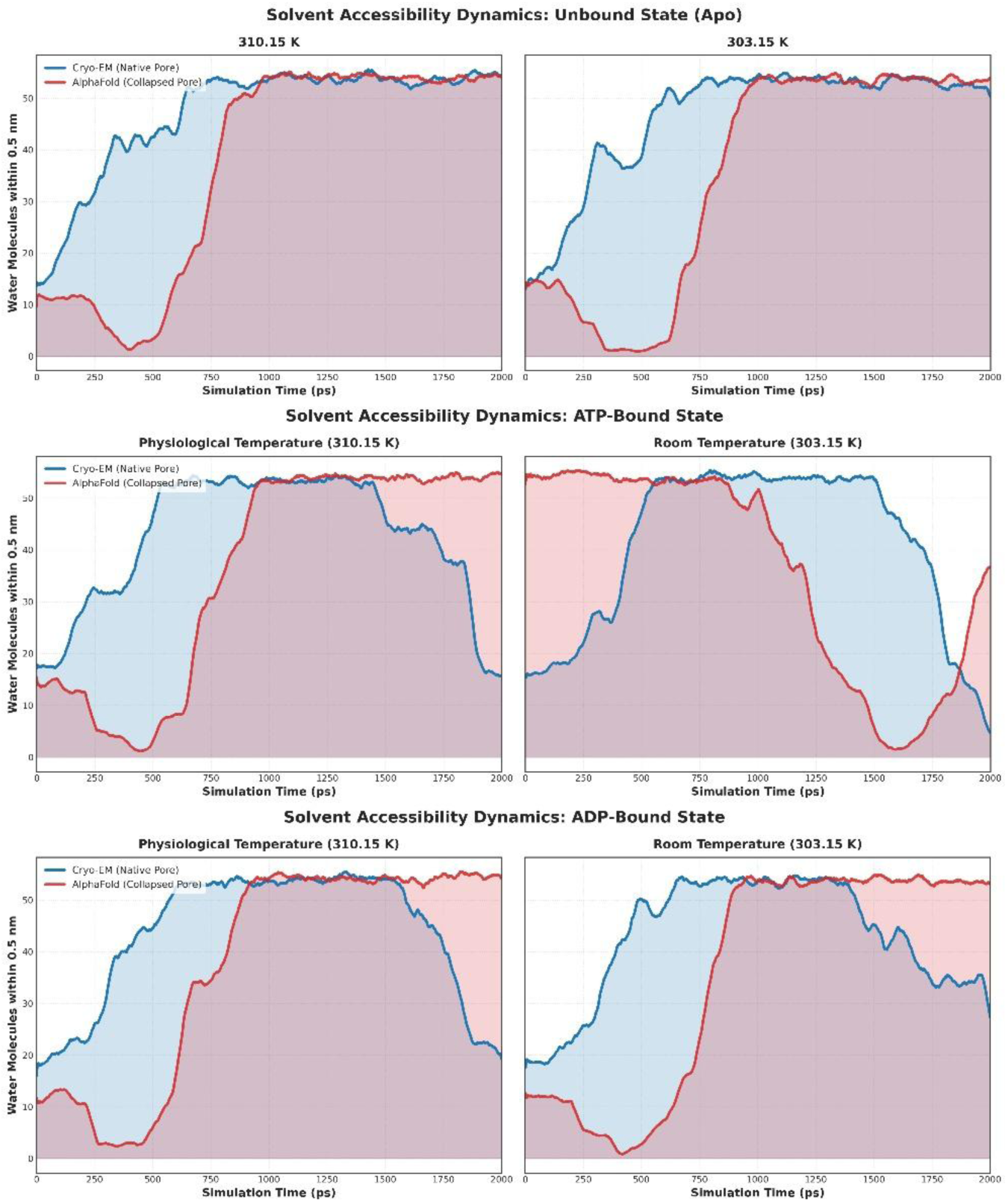
Solvent accessibility dynamics of glucose during steered transport in the apo state. Primary hydration shell (water molecules within 0.5 nm) of glucose across SMD trajectories at 310.15 K (left) and 303.15 K (right) for native Cryo-EM (blue) and AlphaFold (red) models. Native Cryo-EM preserves continuous aqueous hydration along an open transport pathway, whereas AlphaFold exhibits severe early dewetting and delayed rehydration caused by structural pore collapse.

In the unliganded Apo state, the native Cryo-EM framework executes a smooth, progressive solvation pathway across both physiological (310.15 K) and reduced (303.15 K) thermal regimes. Initiating at a baseline of 14 water molecules within 0.5 nm of the substrate, the hexose undergoes continuous, step-wise hydration through an intermediate coordination shelf (41–43 waters) before establishing a stable, bulk-saturated aqueous plateau (52–55 waters) that persists continuously across the central lumen. In sharp contrast, the AlphaFold template subjects the substrate to severe pathological dewetting inside the collapsed central pore. Starting at 11–14 waters, the predicted profile plunges precipitously into an extreme desolvation basin of 1–2 water molecules between 350 ps and 600 ps. This complete stripping of aqueous solvation forces the unliganded glucose through the constricted pore without solvent lubrication, before an abrupt, uncoordinated rehydration surge drives the ligand into a bulk solvent plateau (53–55 waters) where it remains trapped through 2000 ps. This deterministic transition from complete dewetting directly into bulk trapping confirms that structural collapse in predicted Apo GLUT4 destroys the amphiphilic microenvironment required for low-barrier transport.

Under active ATP occupancy, the native Cryo-EM framework maintains a highly controlled, biphasic hydration profile. Commencing at a baseline of 16–18 water molecules, the hexose advances through progressive hydration past an intermediate coordination shoulder (28–32 waters) into a fully hydrated, bulk-like channel envelope (52–55 waters) across the central conduction axis. Upon approaching the extracellular exit portal, the substrate executes orderly, controlled exit desolvation, decaying smoothly to 16–20 waters. Conversely, the ATP-bound AlphaFold model exhibits severe pathological desolvation at 310.15 K, dropping rapidly between 200 ps and 400 ps into a catastrophic dewetting minimum of 1–2 waters before jumping abruptly into bulk saturation (53–55 waters). Under thermal attenuation at 303.15 K, AlphaFold displays anomalous inverted hydration kinetics—initiating bulk-hydrated (54–55 waters) for 900 ps before undergoing a delayed, chaotic desolvation cascade that plunges to 1–2 waters near 1600 ps. While native ATP coordination maintains continuous solvation and functional exit desolvation, the computational model subjects the substrate to severe solvation shocks and thermal volatility.

During ADP coordination, the native Cryo-EM framework preserves a well-behaved solvation trajectory at both 310.15 K and 303.15 K. Starting at an initial baseline of 18–19 water molecules, the substrate ascends steadily through progressive hydration past a prominent intermediate shelf (44–50 waters) to establish a broad, bulk-saturated plateau (53–55 waters) through the central lumen, followed by orderly, controlled exit desolvation down to 20–27 waters near the exit portal. In contrast, the ADP-bound AlphaFold template subjects the substrate to immediate, pathological dewetting, plunging past 200 ps into an extreme desolvation minimum of 1–3 waters between 280 ps and 450 ps. Following this complete stripping of solvent lubrication, the predicted trajectory undergoes abrupt rehydration into an unconstrained bulk plateau (53–55 waters) that remains locked through 2000 ps, failing to execute exit desolvation. These profiles demonstrate that native ADP binding maintains continuous aqueous lubrication matching the unliganded state, whereas structural pore collapse in AlphaFold destroys dynamic microhydration during ADP occupancy.

### 2.7. Electrostatic Surface Remodeling

To determine how nucleotide-induced allostery reshapes the local dielectric environment and governs long-range substrate recognition, solvent-accessible electrostatic surface potentials were mapped across the transporter architecture. The spatial distribution of surface charge dictates not only the long-range steering of polar solutes into the permeation pathway but also the dynamic polarity of the intracellular gate, which directly modulates solvent accessibility and the energetic barriers of conformational transitions. The steady-state electrostatic topologies of the experimental Cryo-EM and predicted AlphaFold frameworks were computationally resolved across the unliganded (Apo), ATP-bound, and ADP-bound ensembles under both physiological (**310.15 K**) and thermally attenuated (**303.15 K**) boundary conditions. This comparative mapping isolates the extent to which specific nucleotide coordination triggers functional charge redistribution—particularly within the critical intracellular vestibule and primary gating zones. Consequently, this analysis establishes whether the inward physical collapse observed in the AI-predicted model induces pathological electrostatic scrambling or preserves the native charge topology strictly required for alternate-access hexose translocation.

#### 2.7.1. Native and Predicted Electrostatic Remodeling at Physiological Temperature (310.15 K)

At 310.15 K, solvent-accessible electrostatic surface potential mapping reveals a dynamic, state-dependent allosteric cycle across the experimental Cryo-EM structure (Figure 22A–C). In the unliganded Apo state, the native transporter adopts an inward-open conformation featuring a deeply splayed cytoplasmic portal framed by basic surface potential, while the extracellular loop network forms a shallow apical concavity lined with electronegative potential. Upon ATP binding, the native structure executes a decisive contraction: the cytoplasmic gate pinches shut to occlude the portal, consolidating basic charge density across a compact base while maintaining an acidic boundary along the extracellular loop crest. Conversely, the ADP-bound state returns to a widely splayed inward-open architecture, reaching maximum aperture flanked by dense electropositive charge clusters. This confirms a functional allosteric mechanism where ATP coordination drives gate closure, whereas Apo and ADP states stabilize an open-inward vestibule for substrate exchange.

**Figure 22.**
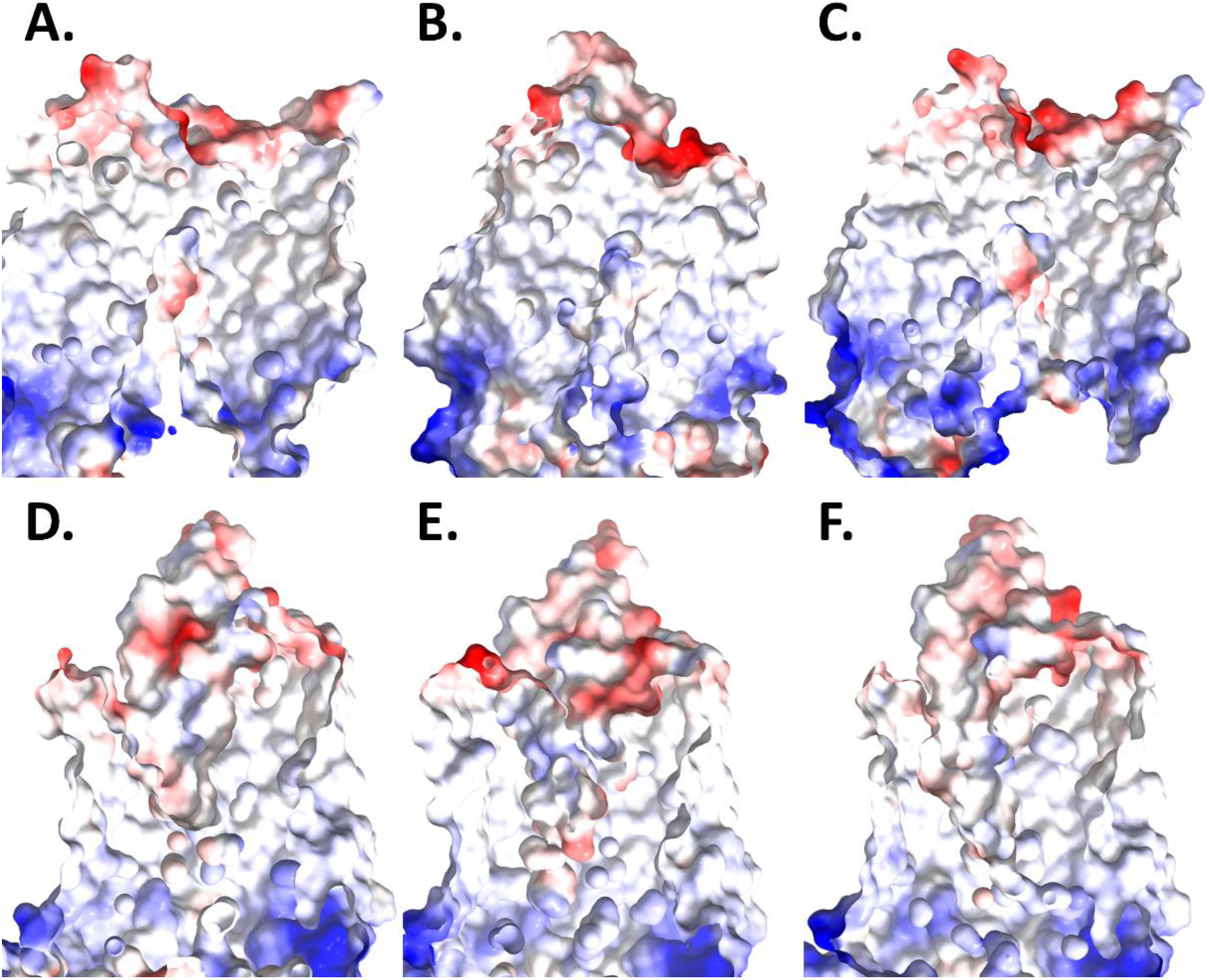
Electrostatic surface remodelling of GLUT4 under physiological conditions (310.15 K). Solvent-accessible electrostatic potential maps (red, negative; blue, positive; white, neutral) for native Cryo-EM **(A–C)** and predicted AlphaFold **(D–F)** structures in apo, ATP-bound, and ADP-bound states.

In contrast, the predicted AlphaFold template remains trapped in a statically constricted, uncoupled fold across all functional states (Figure 22D–F). In the Apo state, the computational model presents a narrowed channel geometry with a prominent, misplaced electronegative patch deeply embedded along the upper-middle luminal wall, interrupting the continuous basic guiding pathway. Under ATP coordination, the model remains pinched at the cytoplasmic base while accumulating an extensive acidic surface region down the upper luminal seam. Crucially, in the ADP-bound state, AlphaFold completely fails to execute cytoplasmic expansion; the portal remains rigidly closed, displaying a fragmented basic rim rather than an open entry funnel.

#### 2.7.2. Temperature-Dependent Conservation of Electrostatic Gating

Under thermal attenuation at 303.15 K, surface potential mapping confirms that state-dependent gating is a thermodynamically resilient property of native GLUT4 (Figure 23G–I). Lower thermal energy preserves the open, basic cytoplasmic portal in the Apo and ADP states, as well as the tightly pinched, electronegative gate region in the ATP-bound complex.

**Figure 23.**
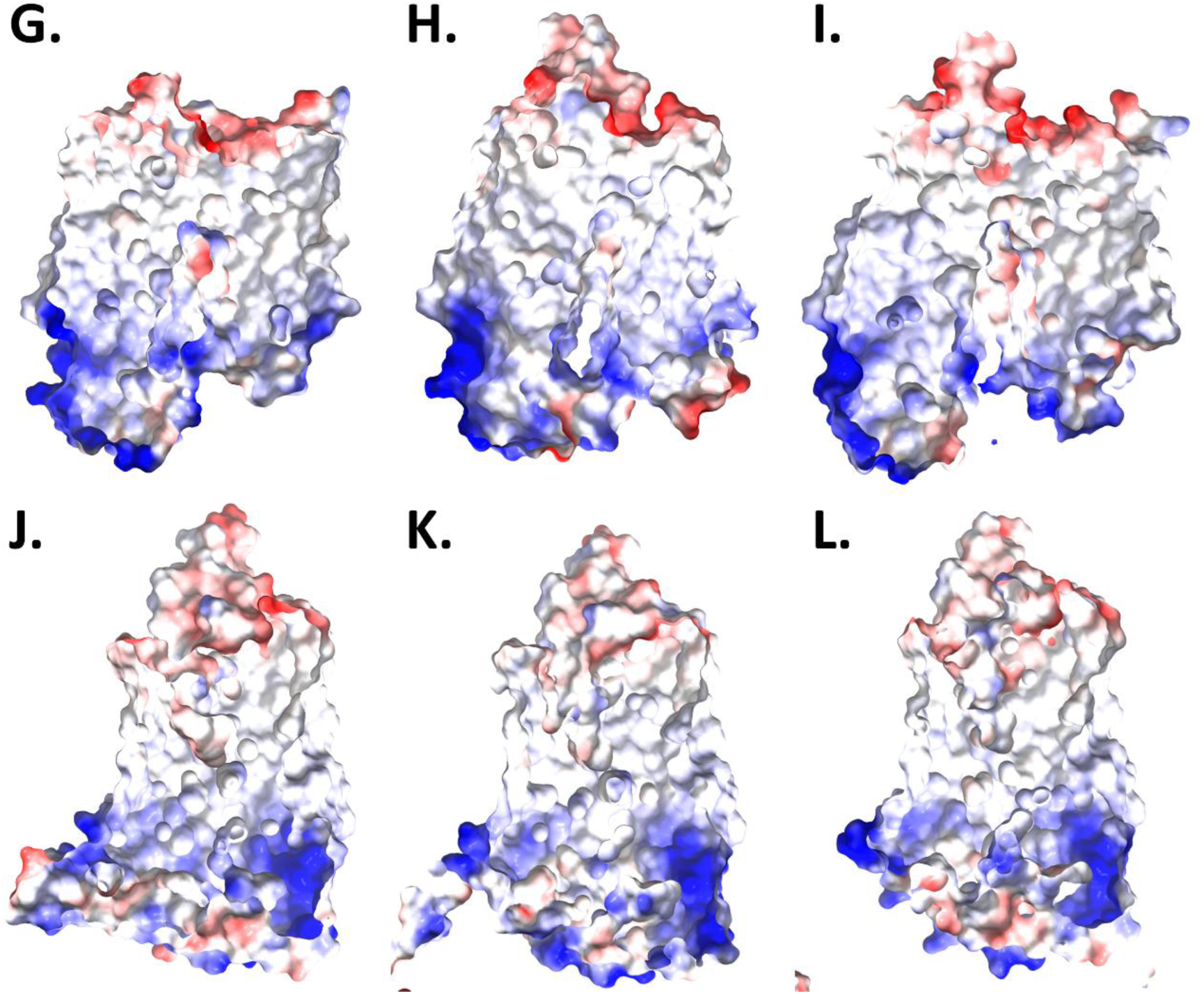
Electrostatic surface remodeling of GLUT4 at 303.15. **K.** Solvent-accessible electrostatic potential maps (red, negative; blue, positive; white, neutral) for native Cryo-EM **(G–I)** and AlphaFold **(J–L)** models in apo, ATP-bound, and ADP-bound states.

Conversely, the AlphaFold template perpetuates its distorted, uncoupled profile across all states at 303.15 K (Figure 23J–L). The model retains a narrowed pore with misplaced luminal acidic patches in Apo, a pinched base in ATP, and a statically closed cytoplasmic gate during ADP occupancy. This persistent charge scrambling and physical constriction across both thermal regimes establish that the AI-predicted structure suffers from a permanent, temperature-invariant flaw that disables alternate-access transport.

### 2.8. Dynamic Cross-Correlation Analysis

Residue-wise dynamic cross-correlation matrices (*C*_*ij*_) constructed from equilibrium molecular dynamics quantify time-averaged spatial covariances between C_*α*_ atom pairs, resolving synchronized correlated (*C*_*ij*_ > 0) and anti-correlated (*C*_*ij*_ < 0) domain fluctuations that govern long-range allosteric communication in GLUT4.

#### 2.8.1. Apo State

At **310.15 K**, unliganded cross-correlation profiles reveal fundamental differences in inter-domain mobility between structural templates (Figure 24). The native Cryo-EM framework displays a balanced, elastic dynamic matrix: strong positive correlations (*C*_*ij*_ ≥ 0.6) remain strictly localized along the main diagonal (intra-helical contacts), while long-range motions between N-terminal (residues 1–230) and C-terminal (residues 270–460) domains exhibit moderate, diffuse anti-correlations (*C*_*ij*_ ≤ −0.3). This unconstrained profile preserves dynamic elasticity for fluid inter-domain tilting. Conversely, the AlphaFold model exhibits severe dynamic hyper-coupling, dominated by massive off-diagonal correlation blocks (*C*_*ij*_ ≥ 0.6) artificially linking distant domains (residues 100–220 and 350–450), interspersed with deep anti-correlation basins (*C*_*ij*_ ≤ −0.5). Thermal attenuation (**303.15 K**) slightly sharpens native diagonal contacts while preserving bounded anti-correlations, whereas AlphaFold perpetuates its hyper-correlated checkerboard layout across all 500 residues, confirming that predicted pore collapse permanently locks the transporter in a rigid state.

**Figure 24.**
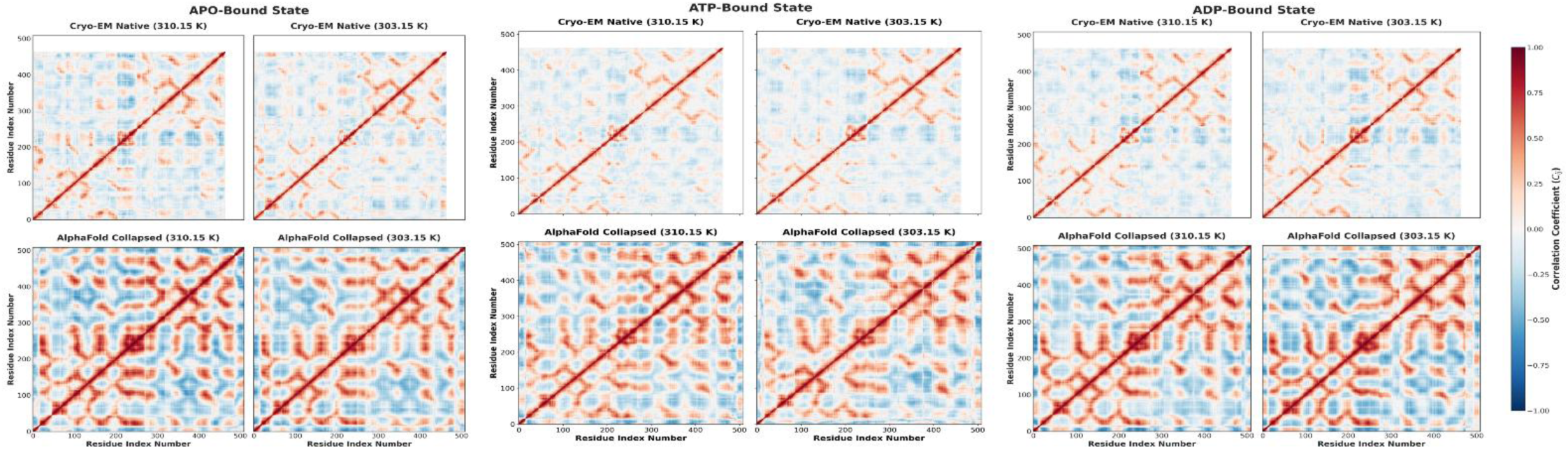
Dynamic Cross-Correlation Matrices of Apo, ATp and ADp bound states of GLUT4. Positive correlation coefficients (red) indicate concerted residue motions, whereas negative correlation coefficients (blue) represent anti-correlated motions. Color intensity reflects the magnitude of the correlation coefficient.

#### 2.8.2. ATP-Bound State

Under physiological ATP coordination (**310.15 K**), the native Cryo-EM framework maintains a fluid cross-correlation profile (Figure 25). Strong diagonal correlations (*C*_*ij*_ ≥ 0.6) capture localized tertiary contacts, while off-diagonal inter-domain regions display gentle, highly diffuse anti-correlations (*C*_*ij*_ ≤ −0.3). This elasticity permits unhindered dynamic sampling between domain halves during nucleotide occupancy. In contrast, the ATP-bound AlphaFold model manifests severe structural lock-up and dynamic frustration, generated by dense off-diagonal correlation blocks (*C*_*ij*_ ≥ 0.6) coupling residues 100–250 with 350–480 alongside deep anti-correlation basins (*C*_*ij*_ ≤ −0.5). Thermal reduction (**303.15 K**) preserves resilient dynamic elasticity in native GLUT4 but fails to relieve AlphaFold’s anomalous grid layout. Persistent off-diagonal hyper-coupling across both temperatures confirms that ATP binding inside the predicted fold traps the protein in an allosterically impaired, hyper-rigid dynamic state.

#### 2.8.3. ADP-Bound State

During ADP occupancy at **310.15 K**, the native Cryo-EM structure exhibits a well-behaved, elastic correlation matrix (Figure 26). Positive correlations (*C*_*ij*_ ≥ 0.6) remain strictly constrained along the main diagonal, while inter-domain regions linking N- and C-terminal halves display pale, diffuse anti-correlations (*C*_*ij*_ ≤ −0.3), preserving fluid domain mobility following loss of the terminal *γ*-phosphate. Conversely, the ADP-bound AlphaFold model perpetuates severe dynamic hyper-coupling, dominated by structured off-diagonal correlation blocks (*C*_*ij*_ ≥ 0.6) that artificially couple residues 100–250 with 350–480, interspersed with exaggerated anti-correlations (*C*_*ij*_ ≤ −0.5). At **303.15 K**, native GLUT4 preserves its elastic communication baseline, whereas AlphaFold fully replicates its rigid checkerboard layout. This temperature-invariant hyper-coupling establishes that structural pore collapse in the computational fold permanently impairs long-range allosteric signaling during ADP coordination.

### 2.9. Dynamic Network Analysis

To translate pairwise correlation patterns derived from dynamic cross-correlation matrices into explicit physical routes of conformational signal transmission, graph-theoretic residue interaction networks (RINs) were constructed. Within these networks, individual C_*α*_atoms function as topological nodes connected by inter-residue edges weighted inversely to pairwise dynamic correlations (*C*_*ij*_). By calculating optimal and suboptimal shortest communication paths between the nucleotide-binding pocket and the intracellular gating region, this analysis isolates critical relay hubs, bottleneck residues, and signal dissipation pathways across unliganded (Apo), ATP-bound, and ADP-bound ensembles under physiological (310.15 K) and thermally attenuated (303.15 K) conditions.

#### 2.9.1. Global Architecture of the Allosteric Communication Network

Shortest-path analysis reveals fundamental topological discrepancies in allosteric signal propagation between the experimental Cryo-EM framework and the predicted AlphaFold model (Figure 27, Table 1). In the native Cryo-EM structure, dynamic allosteric communication operates via an exceptionally conserved, streamlined pathway spanning an invariant topological path length of **4 residues** across all functional states and thermal regimes. At 310.15 K under ATP occupancy, mechanical signals originate at the nucleotide-binding anchor THR331 (THR351), propagate through the histidine relay HSD333 (HSD353), engage the central proline hinge PRO379 (PRO399), and directly terminate at the primary pore-gating residue GLN278 (GLN298) (Figure 27A). This compact 4-step architecture is systematically preserved across ADP-bound and unliganded Apo ensembles at both 310.15 K and 303.15 K (Table 1). The continuous recruitment of the rigid proline hinge (PRO379/PRO399) provides a low-barrier mechanical lever that efficiently couples nucleotide recognition to intracellular gate regulation without signal dissipation.

**Figure 27.**
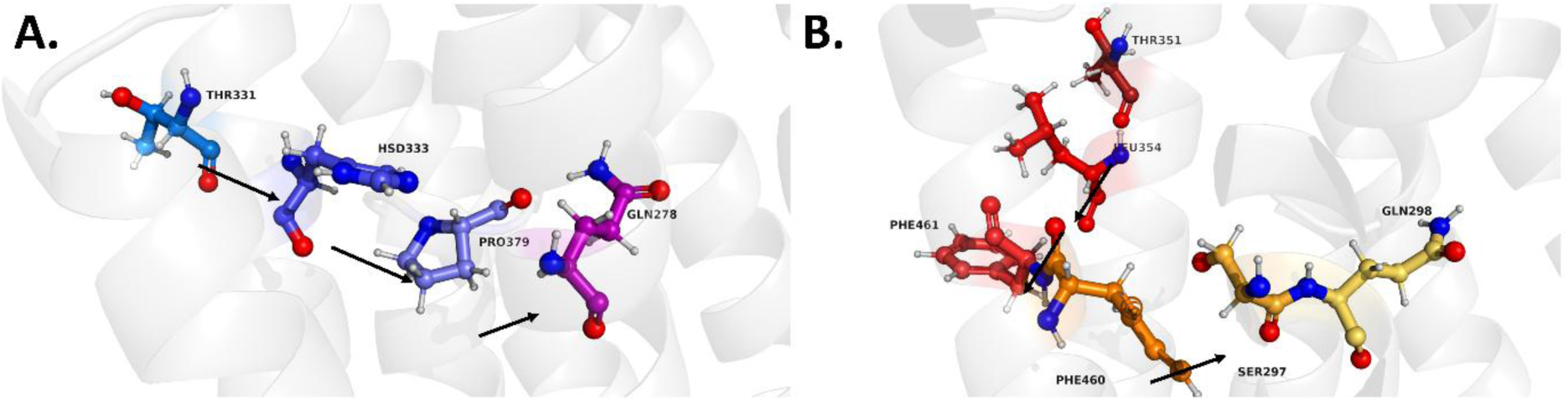
Dynamic allosteric communication pathways in ATP-bound GLUT4 at 310.15 K. **(A)** Cryo-EM structure showing a compact shortest communication pathway from the nucleotide-binding site to the intracellular gate via the native PRO379 hinge. **(B)** AlphaFold structure displaying an elongated pathway rerouted through PHE461–PHE460–SER297, indicating altered network topology. Arrows denote the direction of signal propagation.

**Table 1.** Shortest allosteric communication pathways linking the nucleotide-binding region to the intracellular pore gate.

| System | Model | Ligand | Temperature (K) | Path Length | Shortest Communication Pathway |
| --- | --- | --- | --- | --- | --- |
| A-ATP-310 | AlphaFold | ATP | 310 | 6 | THR351 → LEU354 → PHE461 → PHE460 → SER297 → GLN298 |
| A-ATP-303 | AlphaFold | ATP | 303 | 6 | THR351 → LEU354 → PHE461 → PHE460 → SER297 → GLN298 |
| A-ADP-310 | AlphaFold | ADP | 310 | 6 | THR351 → LEU354 → PHE461 → PHE460 → SER297 → GLN298 |
| A-ADP-303 | AlphaFold | ADP | 303 | 6 | THR351 → LEU354 → PHE461 → PHE460 → SER297 → GLN298 |
| A-APO-310 | AlphaFold | Apo | 310 | 6 | THR351 → LEU354 → PHE461 → PHE460 → SER297 → GLN298 |
| A-APO-303 | AlphaFold | Apo | 303 | 4 | THR351 → LEU352 → PRO399 → GLN298 |
| C-ATP-310* | Cryo-EM | ATP | 310 | 4 | THR331 → HSD333 → PRO379 → GLN278 |
| C-ATP-303* | Cryo-EM | ATP | 303 | 4 | THR331 → HSD333 → PRO379 → GLN278 |
| C-ADP-310 | Cryo-EM | ADP | 310 | 4 | THR351 → HSD353 → PRO399 → GLN298 |
| C-ADP-303 | Cryo-EM | ADP | 303 | 4 | THR351 → HSD353 → PRO399 → GLN298 |
| C-APO-310 | Cryo-EM | Apo | 310 | 4 | THR351 → HSD353 → PRO399 → GLN298 |
| C-APO-303 | Cryo-EM | Apo | 303 | 4 | THR351 → HSD353 → PRO399 → GLN298 |

In sharp contrast, the AlphaFold template exhibits severe topological rewiring and allosteric uncoupling, characterized by a substantially elongated communication pathway spanning a path length of **6 residues** across nearly all simulated conditions (A-ATP-310, A-ATP-303, A-ADP-310, A-ADP-303, and A-APO-310) (Table 1). In the ATP-bound predicted model at 310.15 K, signal transmission originates at THR351 but completely bypasses the native proline hinge. Instead, the signal routes through LEU354, undergoes redirection through a hydrophobic aromatic relay comprising PHE461 and PHE460, passes a polar transfer step at SER297, and finally reaches the gating residue GLN298 (Figure 27B). Only under unliganded thermal attenuation (A-APO-303) does the computational fold shorten to a 4-residue path, though it still utilizes an altered aliphatic bridge (LEU352) rather than the native histidine relay.

#### 2.9.2. Conserved Relay Residues and Communication Hubs

Across all ten simulated structural ensembles, dynamic network analysis identifies two universal terminal anchors that define the physical boundary conditions of allosteric signaling (Table 2). Signals consistently originate at THR351 (THR331) (frequency = 10/10), which functions as the universal nucleotide-binding anchor, and terminate at GLN298 (GLN278) (frequency = 10/10), serving as the universal intracellular pore-gating node. Within the native Cryo-EM framework, signal transfer between these terminal boundary anchors relies on a highly conserved, low-barrier relay module where HSD353 (HSD333) (frequency = 4/6 native ensembles) transfers mechanical torque directly to the central proline hinge PRO399 (PRO379) (frequency = 5/6 native ensembles).

**Table 2.**
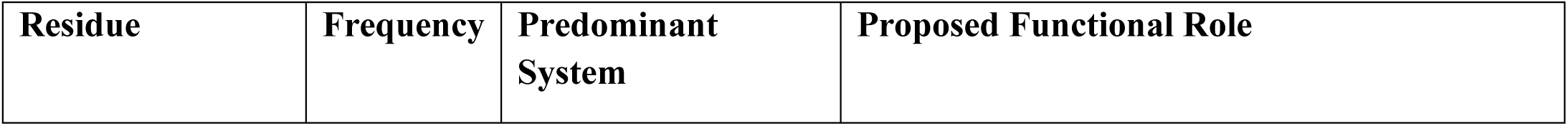
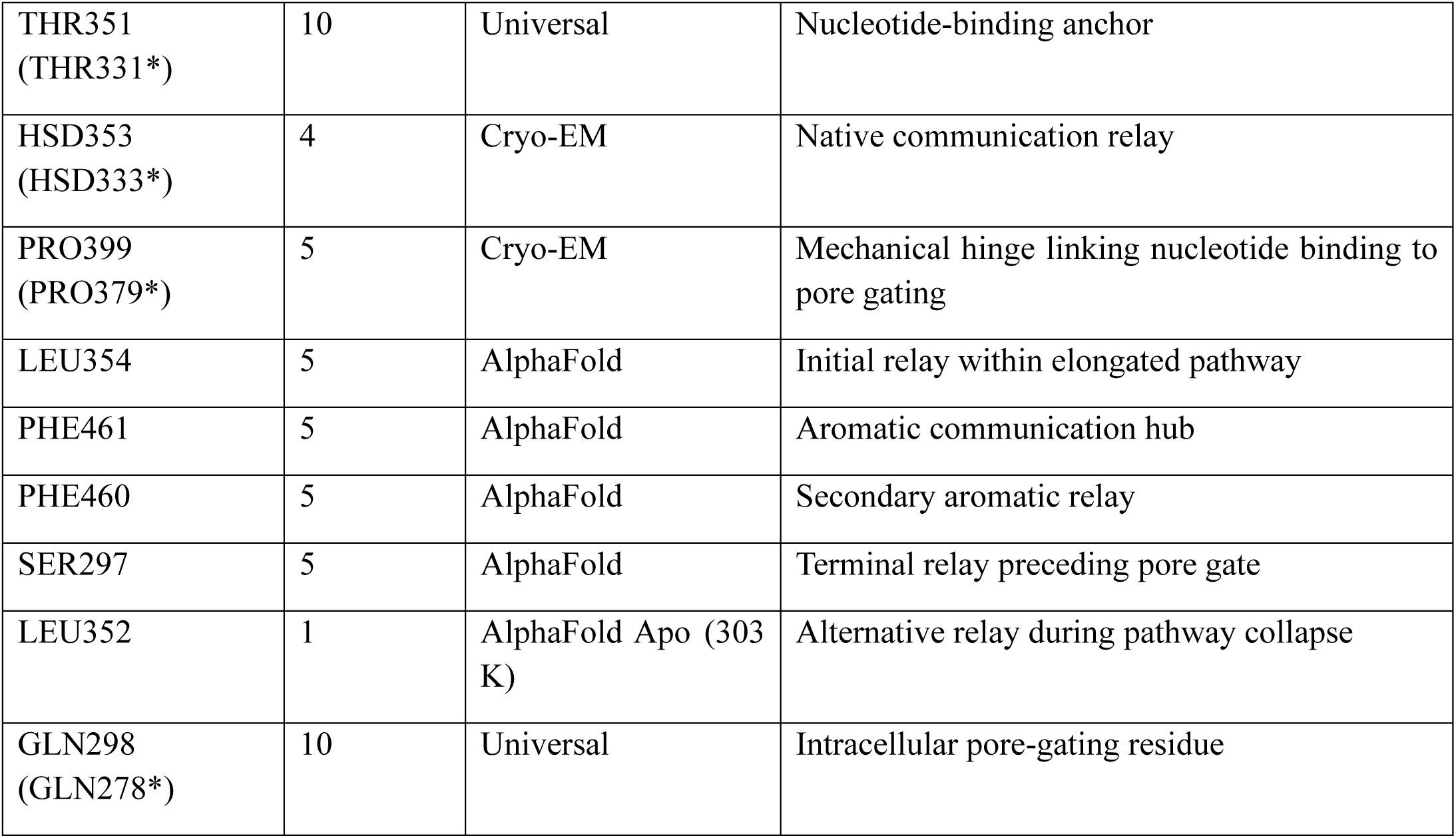
Conserved relay residues identified across dynamic communication pathways.

| Residue | Frequency | Predominant System | Proposed Functional Role |
| --- | --- | --- | --- |
| THR351<br>(THR331*) | 10 | Universal | Nucleotide-binding anchor |
| HSD353<br>(HSD333*) | 4 | Cryo-EM | Native communication relay |
| PRO399<br>(PRO379*) | 5 | Cryo-EM | Mechanical hinge linking nucleotide binding to pore gating |
| LEU354 | 5 | AlphaFold | Initial relay within elongated pathway |
| PHE461 | 5 | AlphaFold | Aromatic communication hub |
| PHE460 | 5 | AlphaFold | Secondary aromatic relay |
| SER297 | 5 | AlphaFold | Terminal relay preceding pore gate |
| LEU352 | 1 | AlphaFold Apo (303 K) | Alternative relay during pathway collapse |
| GLN298<br>(GLN278*) | 10 | Universal | Intracellular pore-gating residue |

Conversely, structural distortion in the AlphaFold model completely dismantles this native signaling axis, routing allosteric signals through a bulky, non-native relay network (Table 2). In the predicted fold, signals bypass the native histidine-proline axis, initiating through LEU354 (frequency = 5/6 AF ensembles) as the primary aliphatic relay. The signal then enters a rigid aromatic communication hub consisting of PHE461 (frequency = 5) and its adjacent secondary relay PHE460 (frequency = 5). From this hydrophobic stack, mechanical strain is transmitted through SER297 (frequency = 5) as a terminal polar relay preceding GLN298. Replacing the streamlined native proline lever with an extended PHE461--PHE460 aromatic stack introduces significant conformational drag, confirming that structural pore collapse in AlphaFold replaces an efficient mechanical coupling with an energy-dissipative pathway.

#### 2.9.3. Network Conservation Across Functional States

Comparative evaluation of dynamic network parameters underscores the stark contrast in allosteric stability between the two structural models (Table 3). The experimental Cryo-EM framework exhibits absolute topological conservation across all functional ligand states (Apo, ATP, ADP) and thermal regimes (310.15 K and 303.15 K). Native GLUT4 maintains an invariant 4-residue pathway length (THR351 → HSD353 → PRO399 → GLN298) with negligible temperature dependence, preserving a robust, low-friction mechanical link between nucleotide recognition and intracellular gate opening.

**Table 3.** Comparative characteristics of dynamic communication networks in Cryo-EM and AlphaFold GLUT4 models.

| Feature | Cryo-EM (Native) | AlphaFold (Predicted) |
| --- | --- | --- |
| Dominant pathway length | 4 residues | 6 residues |
| Primary communication route | THR351 → HSD353 → PRO399 → GLN298 | THR351 → LEU354 → PHE461 → PHE460 → SER297 → GLN298 |
| Principal relay residue | PRO399 (PRO379*) | PHE461–PHE460 aromatic pair |
| Network topology | Compact and conserved | Elongated and rerouted |
| ATP vs ADP | Identical pathway | Identical pathway |
| Apo pathway | Conserved | Alternative pathway only at 303 K |
| Temperature dependence | Negligible | Negligible |
| Structural implication | Efficient native mechanical coupling | Extended communication through an alternative relay network |
| Overall interpretation | Stable and conserved allosteric wiring | Distinct communication architecture associated with the predicted structural model |

In contrast, the predicted AlphaFold model establishes a structurally altered communication architecture defined by pathway elongation (dominant 6-residue length) and topological rerouting through the PHE461--PHE460 aromatic pair (Table 3). Although this elongated topology remains identical between predicted ATP and ADP complexes, the unliganded Apo state exhibits transient pathway instability under thermal attenuation (A-APO-303). This rerouting demonstrates that structural collapse in AlphaFold reorganizes the native communication network into an extended relay, impairing long-range allosteric coupling efficiency across the transporter matrix.

## 3. Discussion

### 3.1. Mechanistic Synthesis: Dynamic Plasticity vs. Static Trapping

Determining the structural and thermodynamic factors that differentiate transport-competent states from statically trapped conformations is essential for understanding facilitated glucose diffusion. Comparative evaluation across twelve distinct structural ensembles demonstrates that while AI architectures accurately predict global fold topology, they can fail to reproduce the dynamic plasticity required for functional transport. Across extended molecular dynamics simulations, the native Cryo-EM framework maintains structural integrity, rapidly establishing an asymptotic backbone plateau (**RMSD = 0.08–0.13 nm**) across unliganded (Apo), ATP-bound, and ADP-bound states under both physiological (**310.15 K**) and thermally attenuated (**303.15 K**) conditions. In contrast, the AlphaFold model undergoes continuous structural drift (**terminal RMSD = 0.33–0.50 nm**). This macroscale divergence demonstrates that transport function depends on dynamic boundary containment and structural elasticity rather than static backbone topology alone.

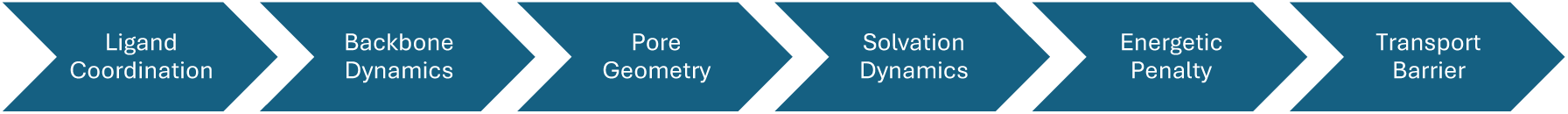

### 3.2. Structural Dynamics and Pore Architecture Dislocation

This dynamic instability arises from a direct causal chain linking local nucleotide coordination, backbone packing, and pore geometry:

- **Local Nucleotide Coordination:** Native ATP binding assembles a multivalent coordination nest (**ARG447**, **ARG462**, **PHE445**) around the triphosphate tail, stabilizing the core translocation domain (residues 230–280; **RMSF = 0.04–0.09 nm**). AlphaFold relies on sparse contacts (**ARG472**, **ARG474**), leaving the nucleoside unoptimized and inducing local core volatility (**RMSF = 0.15–0.19 nm**).
- **Intramolecular Hydrogen Bonding:** This loss of flexibility drives internal over-packing, with the AlphaFold template maintaining an unphysiological surplus of **+15 to +25** intramolecular hydrogen bonds relative to Cryo-EM.
- **Pore Architecture Dislocation:** Spatial radius profiling reveals that while Cryo-EM maintains an open lumen with localized cytoplasmic gating (**bottleneck radius = 0.05 nm**), AlphaFold shifts its primary physical bottleneck extracellularly into the central lumen, dropping below the **0.14 nm** steric threshold required for pore hydration.

### 3.3. Solvation Dynamics and Transport Energetics

This central luminal constriction alters the aqueous microenvironment required for substrate translocation. Steered molecular dynamics (SMD) simulations show that hexose transport through amphiphilic channels requires continuous water-mediated lubrication:

- **Microhydration Shell Dynamics:** The Cryo-EM structure preserves a well-hydrated pathway, guiding glucose through intermediate coordination shelves (**41–50 waters**) into a bulk-saturated envelope (**52–55 waters**) before orderly exit desolvation (**16–20 waters**). AlphaFold’s central bottleneck causes catastrophic early dewetting, stripping the primary shell down to **1–2 waters**.
- **Interaction Energy Penalties:** Forcing unlubricated glucose through the collapsed predicted pore traps the substrate in deep Lennard-Jones steric wells (**-95 to -100 kJ/mol**) and intense Coulombic polar entrapment (**-115 to -175 kJ/mol**).
- **Non-Equilibrium Work Barriers:** Consequently, while native Cryo-EM requires minimal cumulative work (*W*_raw_ = 174.4--223.8 kJ/mol), the steric gridlock and desolvation penalties in AlphaFold generate a massive energetic barrier (*W*_raw_ = 476.3--710.9 kJ/mol).

### 3.4. Allosteric Networks, Dynamic Correlation, and Communication Topology

Pore collapse disrupts the long-range allosteric communication networks that couple nucleotide recognition to gating dynamics:

- **Electrostatic Surface Topology:** Native GLUT4 exhibits dynamic charge remodeling (open basic portal in Apo/ADP; compact gate closure in ATP). AlphaFold presents a static, misplaced electronegative patch along the upper luminal wall that interrupts the basic guiding pathway.
- **Dynamic Cross-Correlations (***C*_*ij*_): Cryo-EM maintains balanced covariance with diffuse inter-domain anti-correlations (*C*_*ij*_ ≤ −0.3). AlphaFold displays rigid hyper-coupling (*C*_*ij*_ ≥ 0.6) and deep anti-correlation basins (*C*_*ij*_ ≤ −0.5).
- **Communication Pathways:** Graph-theoretic analysis reveals that native signals traverse a compact **4-residue path** (THR351/331 → HSD353/333 → PRO399/379 → GLN298/278) mediated by an invariant proline hinge. AlphaFold reroutes signaling into an elongated **6-residue path** (THR351 → LEU354 → PHE461 → PHE460 → SER297 → GLN298) through a bulky PHE461--PHE460 aromatic stack.

### 3.5. Biological Implications for GLUT4 Transport Mechanics

The terminal *γ*-phosphate of ATP acts as the primary chemical trigger for allosteric gating. Loss of this group in ADP reduces net charge demand, driving local side-chain relaxation (**ARG472**, **ARG474**, **GLY473**) and resetting GLUT4 to an open-inward conformation.

This allosteric reset explains why ADP-bound GLUT4 maintains a low-friction, hydrated conduction axis (*W*_raw_ = 176.1 kJ/mol; **0.38 nm** exit aperture) nearly identical to the unliganded Apo state (174.4 kJ/mol), enabling unhindered substrate exchange under elevated intracellular ADP concentrations. Conversely, ATP coordination executes controlled, low-barrier gate tightening (223.8 kJ/mol). Furthermore, identifying **PRO399 (PRO379)** as an invariant relay node highlights the role of proline backbone hinges in MFS transporters, where the sterically restricted ring efficiently transmits torque from the nucleotide pocket to the intracellular gate **GLN298 (GLN278)**.

### 3.6. Implications for AI-Driven Structural Biology and MD Simulations

- **Topology vs. Dynamics:** AlphaFold accurately predicts global tertiary architecture. However, because training metrics emphasize static backbone accuracy (C_*α*_-RMSD, lDDT), AI models can collapse into hyper-packed ground states that omit local side-chain flexibility and dynamic water networks.
- **Caution in Virtual Screening:** Directly using static AI predictions for molecular docking, virtual screening, or non-equilibrium transport simulations risks introducing severe artifacts, including artificial steric bottlenecks and false desolvation barriers.
- **The Imperative for Dynamic Validation:** Functional transport depends on dynamic ensembles. Future workflows integrating predictive models must combine machine-learning structures with explicit-solvent MD, solvation profiling, and network analysis to validate translocation pathway dynamics prior to downstream applications.

## 4. Methods

### 4.1. Structural Preparation and Molecular Docking

Comparative MD simulations utilized two human GLUT4 structures: an inward-open Cryo-EM model (PDB: 7WSN) \cite{yuan_cryo-em_2022} and an AlphaFold prediction (UniProt: P14672) \cite{jumper_highly_2021} terminally C-capped (COOH^−^). Structures were processed in UCSF Chimera to remove non-protein atoms. Ligands (glucose, ATP, ADP) converted via OpenBabel \cite{oboyle_open_2011} and Avogadro \cite{hanwell_avogadro_2012} were parameterized for CHARMM36 using CGenFF \cite{vanommeslaeghe_automation_2012}. Site-specific docking in AutoDock Vina \cite{eberhardt_autodock_2021} targeted the transmembrane \texttt{LSQQL} motif for glucose (Site G) \cite{paul_w_hruz_structural_2001} and the cytoplasmic \texttt{GRRTLHL} motif for nucleotides (Site L) \cite{bazuine_genistein_2005}, generating *apo*, ATP-bound, and ADP-bound complexes validated via PLIP \cite{schake_plip_2025} and Maestro \cite{jacobson_automated_2017}.

### 4.2. Membrane Assembly and System Equilibration

Complexes were oriented via PPM 2.0 and embedded into a POPC bilayer (112 lipids/leaflet) using CHARMM-GUI \cite{park_charmm-gui_2026}. Systems were solvated (2.5 nm water buffer), neutralized (0.15 M KCl, 0.2 M CaCl_2_), and simulated using GPU-accelerated GROMACS \cite{abraham_gromacs_2026, hess_gromacs_2008} with the CHARMM36 force field. Integration used a 2 fs time step with LINCS constraining H-bonds \cite{hess_lincs_1997}, PME electrostatics, and a 1.2 nm van der Waals cutoff. Following steepest-descent energy minimization, a 6-stage NPAT equilibration regulated temperature (V-rescale thermostat) and normal pressure (C-rescale barostat) under periodic boundary conditions.

### 4.3. Steered Molecular Dynamics

Constant-velocity SMD (cv-SMD) probed hexose translocation energetics across *apo*, ATP-bound, and ADP-bound states under physiological (310.15 K) and reduced (303.15 K) temperatures. To ensure statistical reproducibility, each scenario across both Cryo-EM and AlphaFold frameworks was executed in independent triplicates (*n* = 3). Glucose was pulled along the membrane normal at 0.25 nm/ps using a virtual harmonic spring (*k* = 500 kJ/mol/nm^2^). Applied force and displacement coordinates were continuously recorded to compute cumulative raw mechanical work (*W*_raw_) profiles and ensemble standard deviations.

### 4.4. Work Profiles

To evaluate the energetic cost of substrate translocation, constant-velocity Steered Molecular Dynamics (cv-SMD) simulations pulled the glucose substrate through the central conduction pore along the membrane normal (*Z*-axis). Raw cumulative mechanical work (*W*_raw_) was computed by numerically integrating the instantaneous pulling force (*F*_ligand_) with respect to displacement distance (*dx*) (Park & Schulten, 2004):

#### 1. Equation (60) — Section III.A (Page 5953)

This is the main definition of cumulative work during a pulling simulation:

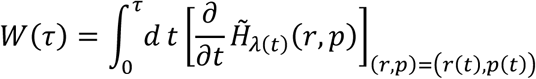

#### How this translates to *W* = ∫ *F* ⋅ *dx*

In Steered Molecular Dynamics, the time-dependent part of the Hamiltonian (*H̃*) is the harmonic spring potential holding the ligand:

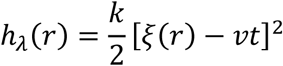

When you take the partial derivative with respect to time *t*:

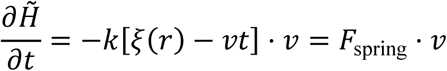

Since 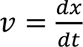 (velocity is displacement over time), multiplying *v* ⋅ *dt* gives *dx*:

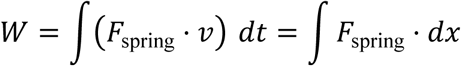

#### 2. Equation (73) — Section III.D (Page 5955)

This is the explicit force-discretized form used for the calculations in their scripts:

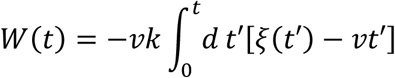

#### Breakdown

- −*k*[*ξ*(*t*′) − *vt*′] = Instantaneous pulling force (*F*_ligand_) exerted by the spring.
- *v dt*′ = Incremental displacement step (*dx*).
- Therefore, −*vk*[*ξ*(*t*′) − *vt*′]*dt*′ = *F*_ligand_ ⋅ *dx*, giving:

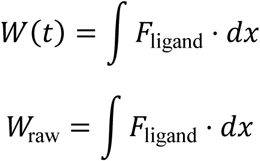

Data processing was executed in Python 3 using NumPy, Pandas, and SciPy. Force (*F*_ligand_) and displacement (*x*) profiles extracted from GROMACS output files (smd_r*.pullf.xvg and smd_r*.pullx.xvg) were interpolated across a standardized common displacement grid (*x* ∈ [0,2.0] nm, 500 points) using scipy.interpolate.interp1d. Cumulative work curves were computed via the trapezoidal integration module scipy.integrate.cumulative_trapezoid. To isolate steric friction and substrate desolvation penalties from background solvent drag and spring fluctuations, substrate-free "Apo" control simulations were performed using identical spring potentials (*k* = 500 kJ/mol/nm^2^) and pulling velocities (0.25 nm/ps). All profiles represent the ensemble mean ± standard deviation across independent triplicates (*n* = 3), rendered using Matplotlib (seaborn-v0_8-whitegrid).

### 4.5. Interaction Energy Decomposition

Non-bonded potential energy landscapes between the GLUT4 matrix (Protein) and glucose (DEX) were decomposed post-simulation. Trajectory coordinate archives (.xtc) were re-evaluated via multithreaded reruns using gmx mdrun -rerun with topology parameter files updated with the energygrps = Protein DEX directive.

Short-range Coulombic (*E*_Coul-SR_) and Lennard-Jones (*E*_LJ-SR_) interaction terms were extracted from binary energy archives (.edr) using the gmx energy utility via automated Python **subprocess** queries targeting specific cross-term matrix indices (Coul-SR:Protein-DEX and LJ-SR:Protein-DEX). Extracted time series were mapped onto a uniform time grid (*t* ∈ [0,2000] ps, 1500 points) using scipy.interpolate.interp1d. To eliminate boundary artifacts without obscuring energetic minima, raw energy timelines were filtered using an edge-preserving moving-average smoothing algorithm (window size = 21 frames) prior to calculating triplicate ensemble statistics.

### 4.6. Pore Geometry Analysis

Spatial radius profiles (*r*) along the central conduction axis (*Z*) were calculated using the **Channel Annotation Package (CHAP)** (Klesse et al., 2019). CHAP quantified luminal cross-sectional dimensions, localized volume fluctuations, and bottleneck constraints across Apo, ATP-bound, and ADP-bound ensembles. The 0.14 nm radius threshold was set as the physical boundary required for pore hydration.

### 4.7. Solvent Accessibility and Hydration Shell Dynamics

Substrate microhydration was monitored across SMD trajectories by tracking water molecules within the primary solvation shell, defined as a 0.5 nm radial boundary surrounding glucose (resname DEX). At 1 ps intervals, water atoms occupying this volume were extracted from coordinate trajectories (smd_r*.xtc) and topology files (smd_r*.tpr) using the GROMACS selection engine gmx select via the Python **subprocess** module:

Raw water atom counts were divided by three to yield discrete, intact TIP3P water molecule counts (*N*_water_). To filter high-frequency thermal noise while preserving macroscopic dewetting transitions, normalized hydration data were smoothed using a 50-frame thermodynamic rolling average.

### 4.8. Electrostatic Surface Remodelling

Solvent-accessible electrostatic surface potential topologies were generated using **UCSF ChimeraX**. Equilibrium PDB snapshots were stripped of non-protein atoms (delete :509) and hydrogenated (addh). Molecular surfaces were generated (surface) and colored according to Coulombic electrostatic potential (coulombic) across a range of −10 to + 10 kcal/mol ⋅ *e* (red, negative/acidic; white, neutral; blue, positive/basic). Views were oriented along the intracellular entrance portal (turn x 90, clip) to resolve gating accessibility.

### 4.9. DCCM

Residue-wise dynamic cross-correlation matrices (*C*_*ij*_) were constructed from equilibrium trajectories to evaluate allosteric coupling between C_*α*_ atoms. To remove rigid-body rotational and translational tumbling, trajectories were fitted to Frame 0 reference structures using gmx trjconv.

The normalized cross-correlation coefficient *C*_*ij*_ was calculated from C_*α*_ displacement vectors (Δ*r*) over 2 ns trajectories (Ichiye & Karplus, 1991):

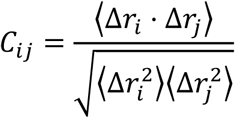

Covariance matrices from independent triplicates (*n* = 3) were ensemble-averaged prior to normalization, where *C*_*ij*_ → +1.0 indicates correlated motion, *C*_*ij*_ → −1.0 indicates anti-correlated motion, and *C*_*ij*_ ≈ 0 indicates uncoupled fluctuations.

### 4.10. Dynamic network Analysis

Graph-theoretic residue interaction networks (RINs) were constructed using **MDAnalysis** and **NetworkX**. C_*α*_ atoms represented graph nodes, with edges defined between residue pairs maintaining an average spatial distance ≤ 8.0 AA (MDAnalysis.analysis.distances). Edge weights (*w*_*ij*_) were calculated from absolute ensemble-averaged cross-correlations (|*C*_*ij*_|):

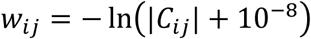

Allosteric signal transmission pathways were mapped using Dijkstra’s shortest-path algorithm (networkx.shortest_path). Boundary nodes were selected based on conserved functional sequence motifs:

- **Source Node (Signal Origin):** THR351 (UniProt register; THR331 in Cryo-EM PDB register), located within the conserved GRRTLHL nucleotide-binding motif (*Carruthers & Helgerson, 1989*).
- **Sink Node (Signal Destination):** GLN298 (UniProt register; GLN278 in Cryo-EM PDB register), located within the conserved QQLS intracellular gating motif on TM7 (*Hruz & Mueckler, 2001*).

## 5. Conclusion

By integrating explicit-solvent molecular dynamics with non-equilibrium steered molecular dynamics and graph-theoretic network analysis, this study resolves the biophysical mechanism of nucleotide-mediated allostery in human GLUT4 while exposing the dynamic limitations of AI-predicted structural models. The native Cryo-EM framework preserves the dynamic plasticity and continuous microhydration necessary for low-barrier hexose transport. ATP coordination acts as an allosteric chemical trigger that tightens the intracellular gate, whereas loss of the terminal *γ*-phosphate in ADP drives local side-chain relaxation, resetting the transporter to an open-inward, low-friction state (*W*_raw_ = 176.1 kJ/mol) nearly identical to the unliganded Apo control. This functional state-dependent gating relies on a robust, topologically conserved four-residue allosteric pathway (THR351 → HSD353 → PRO399 → GLN298) anchored by an invariant proline backbone hinge.

In contrast, the predicted AlphaFold template exhibits severe static over-packing and topological dislocation. By over-optimizing intramolecular contact networks, the computational fold collapses the central permeation axis, inducing catastrophic early dewetting (1–2 water molecules), extreme steric friction, and an unphysiological transport barrier (*W*_raw_ > 470 kJ/mol). Furthermore, structural collapse reroutes allosteric communication into an elongated six-residue pathway through an energy-dissipative aromatic stack (PHE461--PHE460). These findings demonstrate that while machine-learning architectures accurately predict static ground-state folds, they can omit the dynamic volume fluctuations and solvent networks essential for functional transport. Consequently, integrating explicit-solvent MD, microhydration profiling, and dynamic network analysis is imperative to validate AI-generated models prior to downstream virtual screening and drug discovery applications.

## Credit Author Statement

A.L., C.R. and D.A.G. have conceived the interaction hypothesis. A.L., G.S. and S.D. have conceived the computational set-up. G.S., A.L. and S.D. have developed the numerical code and conducted the simulation study. G.S., A.L., S.D. and C.R. have analysed the results. A.L., G.S. and C.R. have developed the theoretical framework. S.D., A.L. and G.S. have drafted this manuscript. All authors have reviewed the manuscript.

## Acknowledgement

One of the authors (A.L.) acknowledges the St Joseph’s Research and Innovation Centre (SJRI) Research Grant vide Grant No.: 2026RDCSJRI031 under which this work was done. A.L. is grateful to Prof. Dr. Ronald J. Mascarenhas and Rev. Dr. Sumeth M.W. Perera for fruitful discussions. A.L. and G.S. are grateful to Mr. Agasthya S.R. for fruitful discussions. A.L. is further thankful to Prof. Dr. K.M. Manoj and Rev. Dr. Roshan Castelino for insightful discussions.

## Data Availability Statement

All program codes and data may be obtained from the corresponding author (A.L.) upon reasonable request. Computational models for GROMACS simulations may be obtained from Lobo, Allen (2026), “Thermodynamic Profiling of Glucose Translocation in POPC embedded GLUT-4 protein models”, Mendeley Data, V1, doi: 10.17632/d6nzsrt9m9.1.

